# eNOS-dependent S-nitrosylation of the NF-κB subunit p65 has neuroprotective effects

**DOI:** 10.1101/2020.02.04.932772

**Authors:** Ariel Caviedes, Barbara Maturana, Katherina Corvalán, Alexander Engler, Felipe Gordillo, Manuel Varas-Godoy, Karl-Heinz Smalla, Luis Federico Batiz, Carlos Lafourcade, Thilo Kaehne, Ursula Wyneken

**Affiliations:** Laboratorio de Neurociencias, Universidad de Los Andes, Santiago, Chile; Institute of Experimental Internal Medicine, Otto-von-Guericke University, Magdeburg, Germany; Universidad Católica del Maule, Chile; Cancer Cell Biology Lab, Centro de Biología Celular y Biomedicina (CEBICEM), Facultad de Medicina y Ciencia, Universidad San Sebastián, Lota 2465, Santiago 7510157, Chile; Leibniz Institute for Neurobiology, Magdeburg, Germany

**Keywords:** NMDA, S-nitrosylation, proteomics

## Abstract

Cell death by glutamate excitotoxicity, mediated by N-methyl-D-aspartate (NMDA) receptors, negatively impacts brain function, including but not limited to hippocampal neurons. The NF-κB transcription factor (composed mainly of p65/p50 subunits) contributes to neuronal death in excitotoxicity, while its inhibition should improve cell survival. Using the biotin switch method, subcellular fractionation, immunofluorescence and luciferase reporter assays, we found that NMDA stimulated NF-κB activity selectively in hippocampal neurons, while endothelial nitric oxide synthase (eNOS), an enzyme expressed in neurons, is involved in the S-nitrosylation of p65 and consequent NF-κB inhibition in cerebrocortical, *i.e*., resistant neurons. The S-nitro proteomes of cortical and hippocampal neurons revealed that different biological processes are regulated by S-nitrosylation in susceptible and resistant neurons, bringing to light that protein S-nitrosylation is a ubiquitous post-translational modification, able to influence a variety of biological processes including the homeostatic inhibition of the NF-κB transcriptional activity in cortical neurons exposed to NMDA receptor overstimulation.

## Introduction

Neuronal death by glutamate excitotoxicity is implicated in the pathogenesis of several neurological disorders, ranging from neurodegeneration to epilepsy, stroke and traumatic brain injury^1,2^. Overstimulation by glutamate leads to massive calcium influx, mainly through N-methyl-D-aspartate receptors (NMDA-Rs), triggering several intracellular pro-death signaling pathways^3^. Endogenous/homeostatic protective mechanisms in response to glutamate, are incompletely known.

In that line, the nuclear factor kappa B (NF-κB) family of transcription factors has been implicated in excitotoxicity in the retina, the striatum, cerebral cortex and hippocampus^4,5,6^. This is associated with induction of pro-apoptotic and pro-inflammatory genes, including IL-1β. The canonical activation of NF-κB depends on phosphorylation and degradation of IκB proteins, leading to release and nuclear translocation of NF-κB, a dimer composed most frequently of a p65 and a p50 subunit^7,8^. Its transcriptional activity in the nucleus is inhibited by S-nitrosylation (*i.e*., the reversible coupling of nitric oxide (NO) to cysteine residues) of the p65 cysteine 38 residue^9,10,11^. However, the contribution of this signaling mechanism to excitotoxicity is unknown. The main source of NO in the brain are nitric oxide synthases, *i.e*., the neuronal (nNOS), endothelial (eNOS) and inducible (iNOS) enzymes^12,13,14^. Considering the novel finding that eNOS is present in neurons and synapses^15^, we examined whether eNOS is involved in p65 S-nitrosylation and, thus, in the regulation of its transcriptional activity under excitotoxicity-promoting conditions. We compared primary cultures of hippocampal and cortical neurons, which differ in their vulnerability to excitotoxic insults: hippocampal neurons have a higher sensitivity than cortical neurons^16^. We found that eNOS contributes to p65 S-nitrosylation and is associated with neuroprotection. This homeostatic mechanism is not active in hippocampal neurons, in which NF-κB activation after an excitotoxic insult leads to increased nuclear translocation and transcriptional activity, including increased transcription of the pro-inflammatory cytokine IL-1β. Our results show that NF-κB activity can be regulated by an eNOS dependent endogenous neuroprotective mechanism in excitotoxicity-like conditions.

## Results

### NF-κB activation in cortical and hippocampal cultures after NMDA stimulation

To assess the participation of NF-κB under excitotoxicity-promoting conditions, we studied the activation and nuclear translocation of p65 in 30 or 100 μM NMDA-stimulated cortical and hippocampal cultures (Figure 1). These cultures contain approximately 30% of astrocytes in addition to neurons^17^. We first assessed cell viability following incubation with different NMDA concentrations (Supplemental Figure S1D): a one hour incubation with any NMDA concentration did not induce cell death in cortical cultures. In turn, in hippocampal cultures, 30 μM NMDA did not induce death while 100 μM NMDA was able to produce substantial cell death when measured 24 hours later. These results are consistent with several reports indicating a time- and concentration dependence of NMDA receptor overstimulation to observe cell death^18,19,20^. We selected 30 μM to 100 μM NMDA for one hour to test the mechanistic steps that participate in the initiation of excitotoxic pathways and that subsequently progress to cell death^21^.

**Figure 1.**
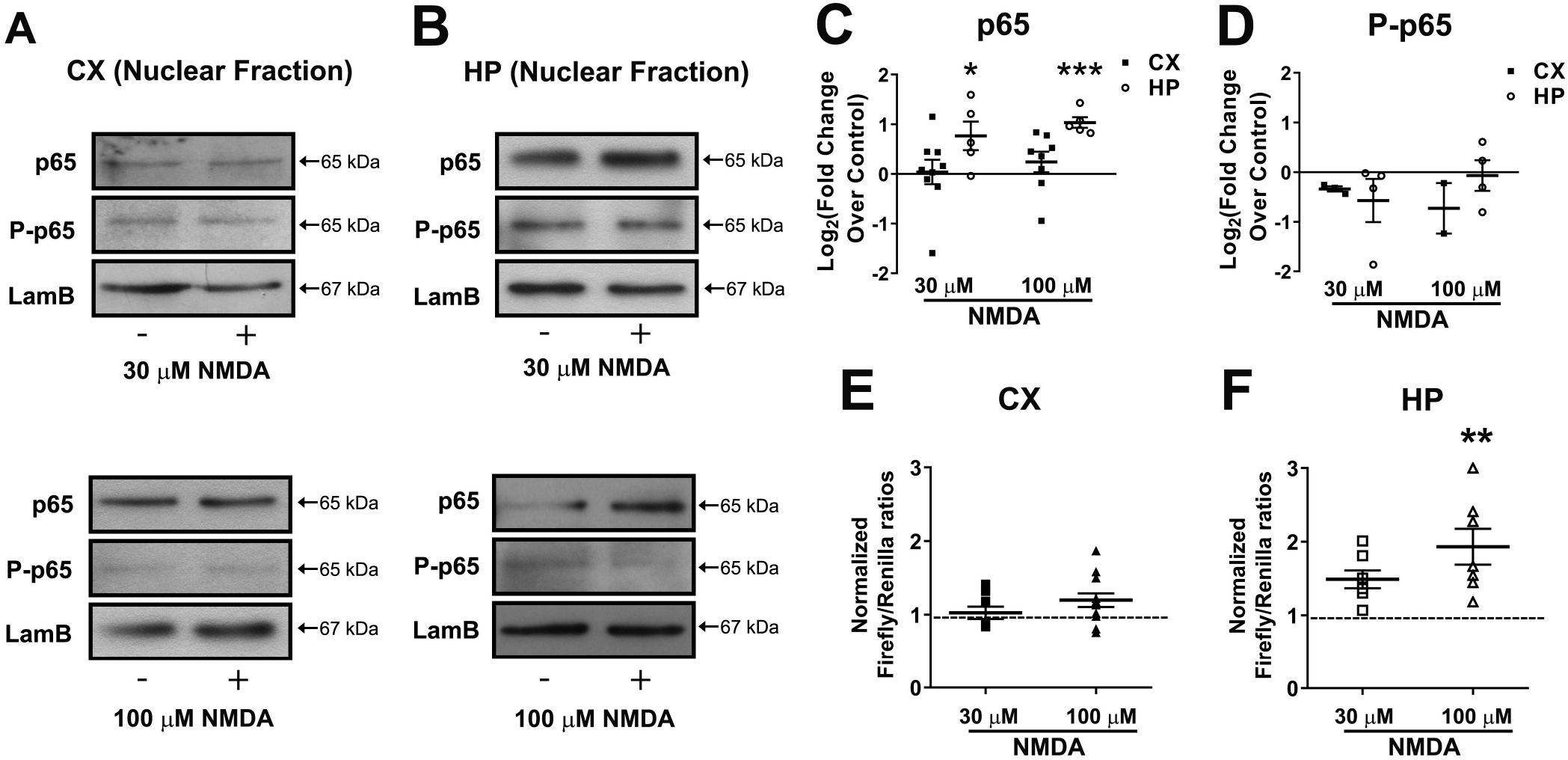
NF-κB is activated in hippocampal, but not in cortical cultures after incubation with NMDA. **A)** and **B)** Neuronal cultures were stimulated with 30 μM or 100 μM NMDA for 1 hour and nuclear fractions were separated subsequently. Representative Western blots of cortical (A) and hippocampal (B) culture-derived nuclear fractions after stimulation with 30 μM (top) or 100 μM (bottom). For each Western blot, equal quantities of proteins were loaded and Lamin B1 (LamB) was used as loading control. **C)** and **D)** Densitometric quantification of relative changes of p65 (C) and phospho-p65 (D) in the nuclear content, comparing stimulated (NMDA) vs control (non-stimulated) condition in the same Western blot. Calculated results obtained of 6 independent experiments (n=6). Statistical significance was assessed by two-tailed t-test (* p<0.05; ** p<0.01). **E)** and **F)** Showing relative luciferase activity in cortical (E) and hippocampal (F) neurons after stimulation with 30 μM or 100 μM NMDA for one hour (n=6). Statistical significance was assessed by One-way ANOVA followed by Bonferroni post-test (**p<0.01).

We first quantified the nuclear translocation of p65. Based on the distribution of a nuclear (*i.e*., Laminin B) and a cytoplasmic (*i.e*., GAPDH) marker, we could conclude that a reliable separation of nuclei from cytoplasm was obtained (Supplemental Figure S1E). In Figures 1A and 1B, representative Western blots of p65 and its phosphorylated form in the nuclear fractions are shown, where Laminin B was used as a loading control. Note that p65 phosphoserine 536 is considered a general marker of NF-κB activation, especially of the canonical pathway^22^. The densitometric analysis of the Western blots (Figures 1C and 1D) confirmed that p65 increased in the nuclear fractions of hippocampal neurons (HP, white bars), but not in cortical neurons (CX, black bars) exposed to the same NMDA concentrations. Interestingly, this was not accompanied by any changes in the levels of phospho-serine 536, indicating that the nuclear translocation of p65 in our experimental model was independent of this phosphorylation site. To determine whether astrocytes contributed to nuclear translocation in the hippocampal cultures, we used immunofluorescence to detect p65 in DAPI-stained nuclei of neurons (labelled with an antibody against microtubule associated protein 2, MAP2) or astrocytes (labelled with an antibody against glial fibrillary associated protein, GFAP) (Supplemental Figure S2). Consistent with the previous observations, we found that 30 μM NMDA induced an increase in the nuclear content of p65 in both neurons and astrocytes in hippocampal cultures (arrows point to cell nuclei in each culture type and experimental condition). No translocation was observed in cortical cell cultures in either cell type.

To evaluate the transcriptional activity of NF-κB, we used the NF-κB luciferase reporter assay (Figure 1E and F). Consistent with the previous results, NF-κB transcriptional activity increased in hippocampal neurons exposed to 100 μM NMDA, while no effects were observed in cortical cells. To test whether NF-κB activation is associated with cell death, we used the NF-κB inhibitor Ro 106-9920 (Figure 2) at a concentration of 2 μM for one hour, not affecting neuronal cell survival *per se* under our experimental conditions (Figure 2A), which is consistent with previous concentration and time-dependence studies using this inhibitor^23,24^. Hippocampal cell death induced by 100 μM NMDA was prevented by NF-κB inhibition with 2 μM Ro106-9920 (Figure 2B-C). Surprisingly, cell death in the cortical cultures (*i.e*., resistant to 100 μM NMDA) increased in the presence of NF-κB inhibition, suggesting opposing roles in neurotoxicity/neuroprotection of NF-κB.

**Figure 2.**
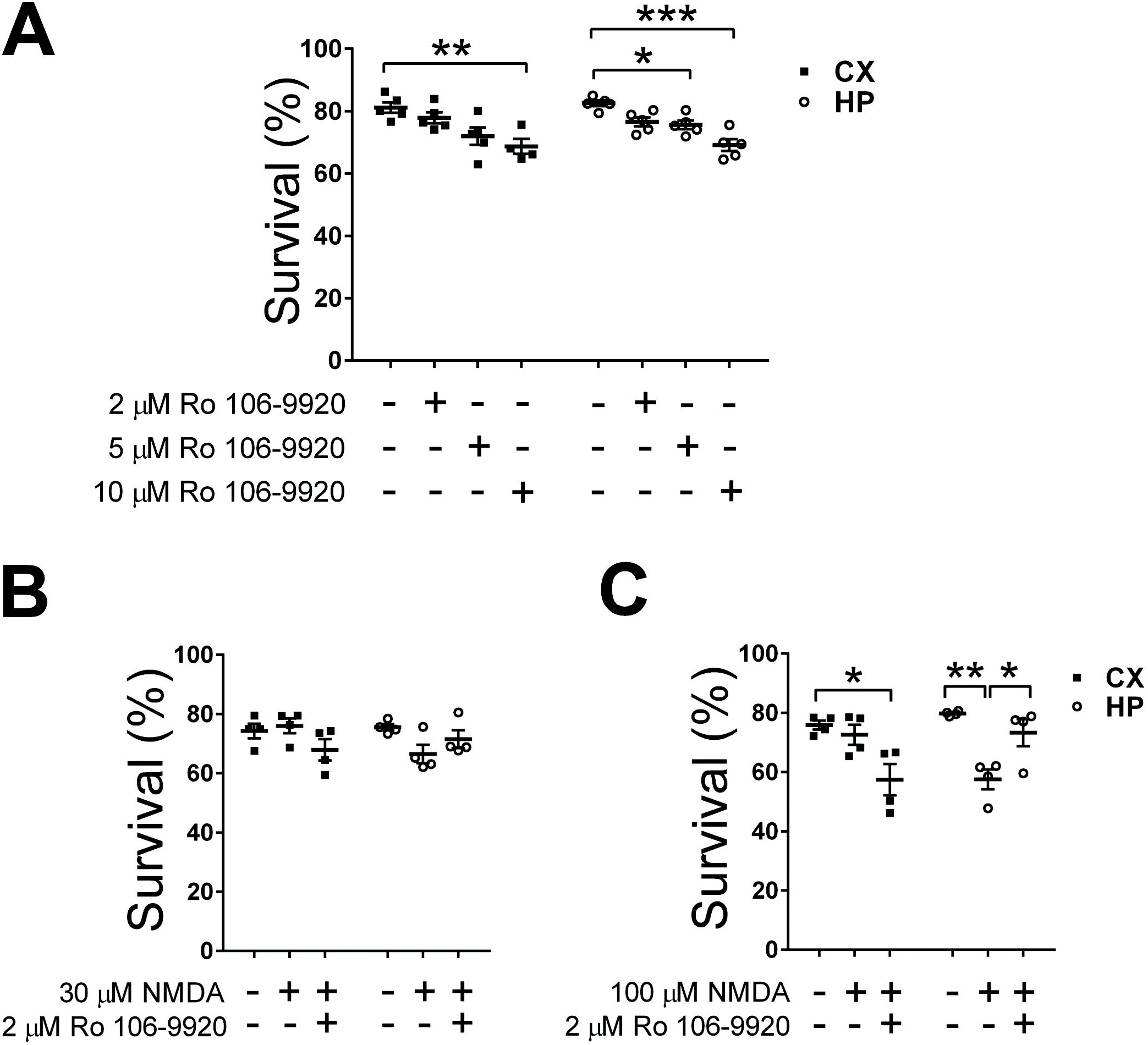
Inhibition of NF-κB with Ro 106-9920 decreases cell viability in cortical cultures but increases it in hippocampal cell cultures. **A)** Effect of different concentrations of 6-(Phenylsulfinyl)tetrazolo[1,5-b]pyridazine (Ro 106-9920) on cell viability of cortical and hippocampal cultures **B) and C)** A concentration of 2 μM Ro-106-9920, chosen because it does not affect cell viability per se, was used in 30 μM (B) or 100 μM (C) NMDA stimulated cultures for one hour. Cell death was detected by Trypan blue exclusion test. Results obtained in n=4 independent experiments. Statistical significance was assessed by One-way ANOVA followed by Bonferroni post-test * p<0.05; ** p<0.01; *** p<0.001.

### S-nitrosylation of p65 increased in cortical cell cultures after NMDA

We then evaluated a potential regulation of the NF-κB p65 subunit by S-nitrosylation using the biotin switch assay^25^. Efficacy of all protocol steps was controlled by Western blot and protein staining (Supplemental Figure S1A-C). Interestingly, the pull down revealed that S-nitrosylation of p65 increased in cortical cells after 30 μM NMDA, while in hippocampal cells the opposite effect was observed (Figure 3). This result supports the idea that regulation of p65 activity by S-nitrosylation is a dynamic post-translational modification. In other experimental models, increased p65 S-nitrosylation is associated with decreased transcriptional activity^9,10,11^.

**Figure 3.**
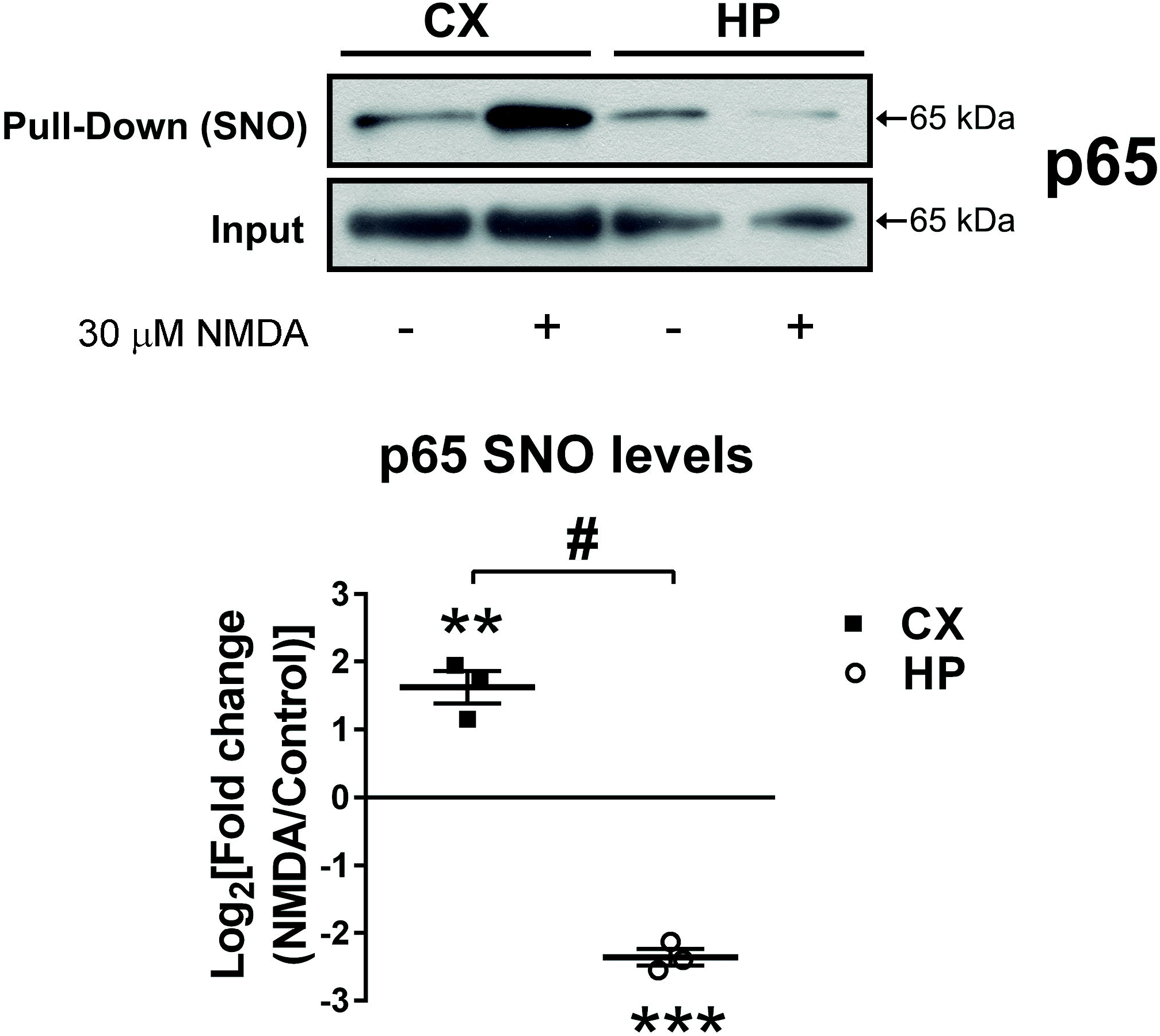
Different levels of NF-κB p65 subunit S-nitrosylation (-SNO) in cortical (CX) and hippocampal (HP) cell cultures after stimulation with NMDA. Neuronal cultures were stimulated with 30 μM NMDA for one hour. Afterwards, cells were homogenized to pull down S-nitrosylated proteins using the biotin switch assay. Representative Western blots of the S-nitosylated p65 subunit of NF-κB and densitometric quantification of cortical and hippocampal cell cultures are shown comparing stimulated (NMDA) vs control (non-stimulated) condition in the same Western blot. n=4 independent experiments and statistical significance was assessed by two-tailed t-test. ** p<0.01; *** p<0.001, # p<0.01.

To evaluate the putative functional effects p65 S-nitrosylation, we directly altered p65 S-nitrosylation by decreasing NO levels by inhibition of nitric oxide synthases (NOS). We focused particularly on eNOS, previously described by us to be expressed in neurons^15^. We measured the eNOS-dependent NO production in cortical cultures transfected with a shRNA targeting eNOS^15^. To stimulate NO production, the neurotrophin BDNF was used^16^. In the presence of the sh-eNOS RNA (but not of a sequence targeting Luciferase as a control), the production of NO decreased, as revealed by the respective slopes (Figure 4A-B).

**Figure 4.**
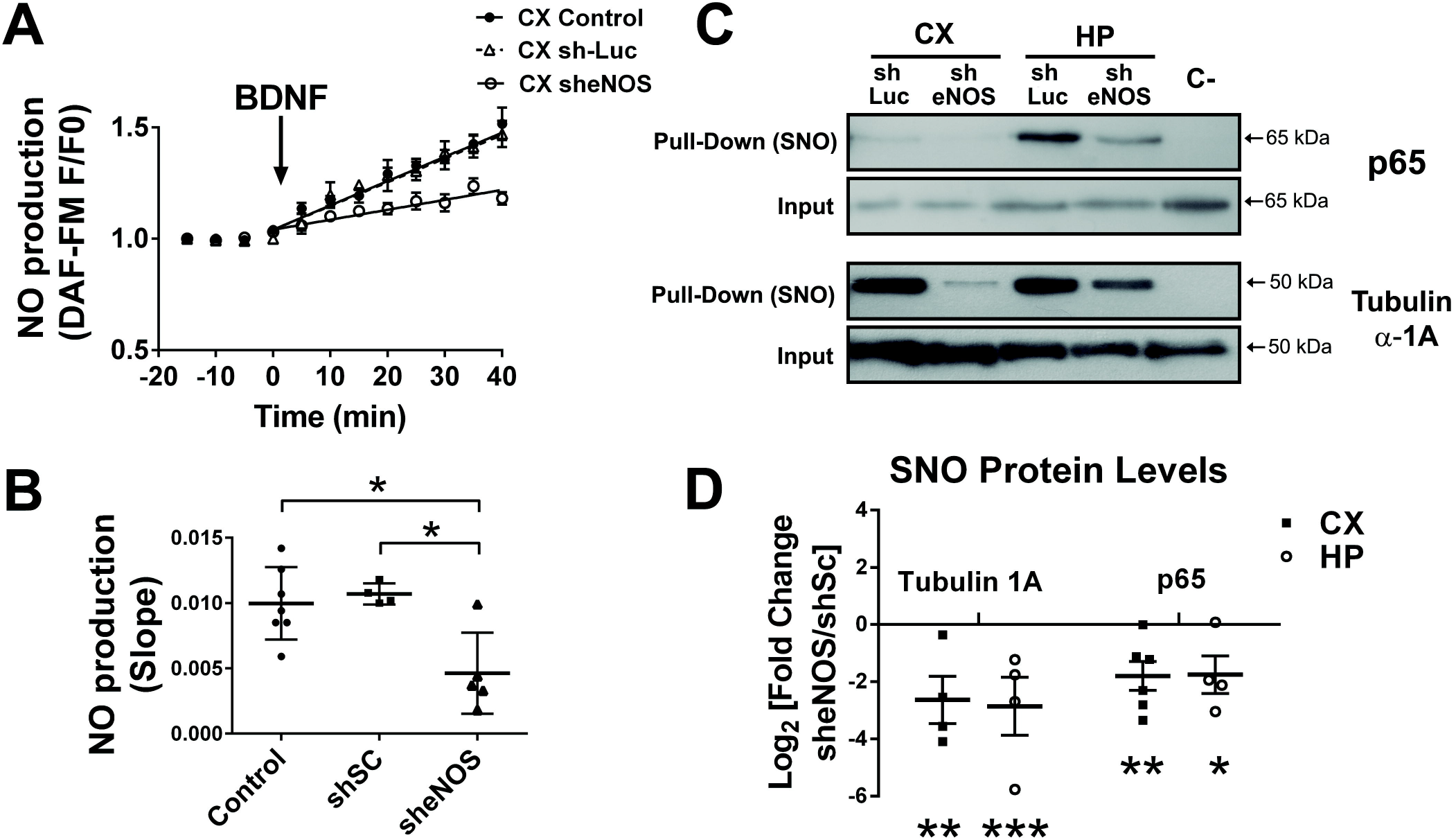
eNOS contributes to NO production and S-nitrosylation of selected proteins. **A)** Relative increase of NO after the addition of 200 ng/ml BDNF to cortical cell cultures previously transfected with a shRNA targeting eNOS, sh-Luc shRNA or not transfected controls. **B)** Mean slopes of NO production are shown in n= 4 to 6 independent experiments, * p<0.5 by two-way ANOVA followed by Bonferroni post-test. **C)** The biotin-switch assay was used to pull down S-nitrosylated proteins. Western blots detecting p65 subunit and tubulin 1α in the pull downs of cortical and hippocampal cultures are shown. Cell cultures were transfected with shRNA targeting eNOS or scrambled shRNA. **D)** Densitometric quantification of the S-nitrosylated (SNO) levels of NF-κB subunit p65 and tubulin α-1A. Result obtained from n= 4 to 6 independent experiments. * p<0.05; ** p<0.01; *** p<0.001 by two-tailed t-test. CX: cortical cultures; HP: hippocampal cultures; Control: not transfected cortical neurons. Sc= scrambled shRNA sequence, eNOS = short interfering RNA against eNOS, C – is a negative control for the biotin switch assay (pull down of samples in which reduction with ascorbate was omitted).

Following, we tested whether decreased endogenous NO production could affect the S-nitrosylation of p65 and tubulin1A, which have been shown to be NO targets^26,27^ (Figure 4CD). In fact, after using the biotin-switch assay of neuronal cultures transfected with sh-eNOS RNA, it was revealed that the S-nitrosylation of p65 and tubulin1A decreased markedly with respect to the sh-Luc controls in cortical and hippocampal cultures. Thus, we conclude that eNOS significantly contributes to the observed protein S-nitrosylation.

### NO regulates transcriptional activity of NF-κB but not its nuclear translocation in response to NMDA stimulation

Although it is known that NO inhibits the transcriptional activity of NF-κB^11^, this type of regulation has not yet been observed in neurons. Moreover, it is unknown whether NO affects nuclear translocation. Therefore, we measured nuclear translocation and NF-κB activity using the general NOS inhibitor LNIO at a concentration of 10 μM (Figure 5)^28,29^. In Figure 5A, we show in nuclear fractionation experiments followed by Western blots that the levels of p65 do not change among the experimental conditions. Moreover, in hippocampal cells the nuclear increase of p65 after 100 μM NMDA could not be prevented by 10 μM LNIO application, suggesting that nuclear translocation is not affected by S-nitrosylation. To evaluate putative changes in p65 expression levels, we compared total p65 within the cellular homogenates. The constant expression levels clearly indicate that the increased nuclear content is a result of enhanced translocation (Supplemental Figure S3A-B). In that line, we also measured the IκB-α levels in the cytoplasm, and we did not find significant differences among groups (Supplemental Figure S3C-D).

**Figure 5.**
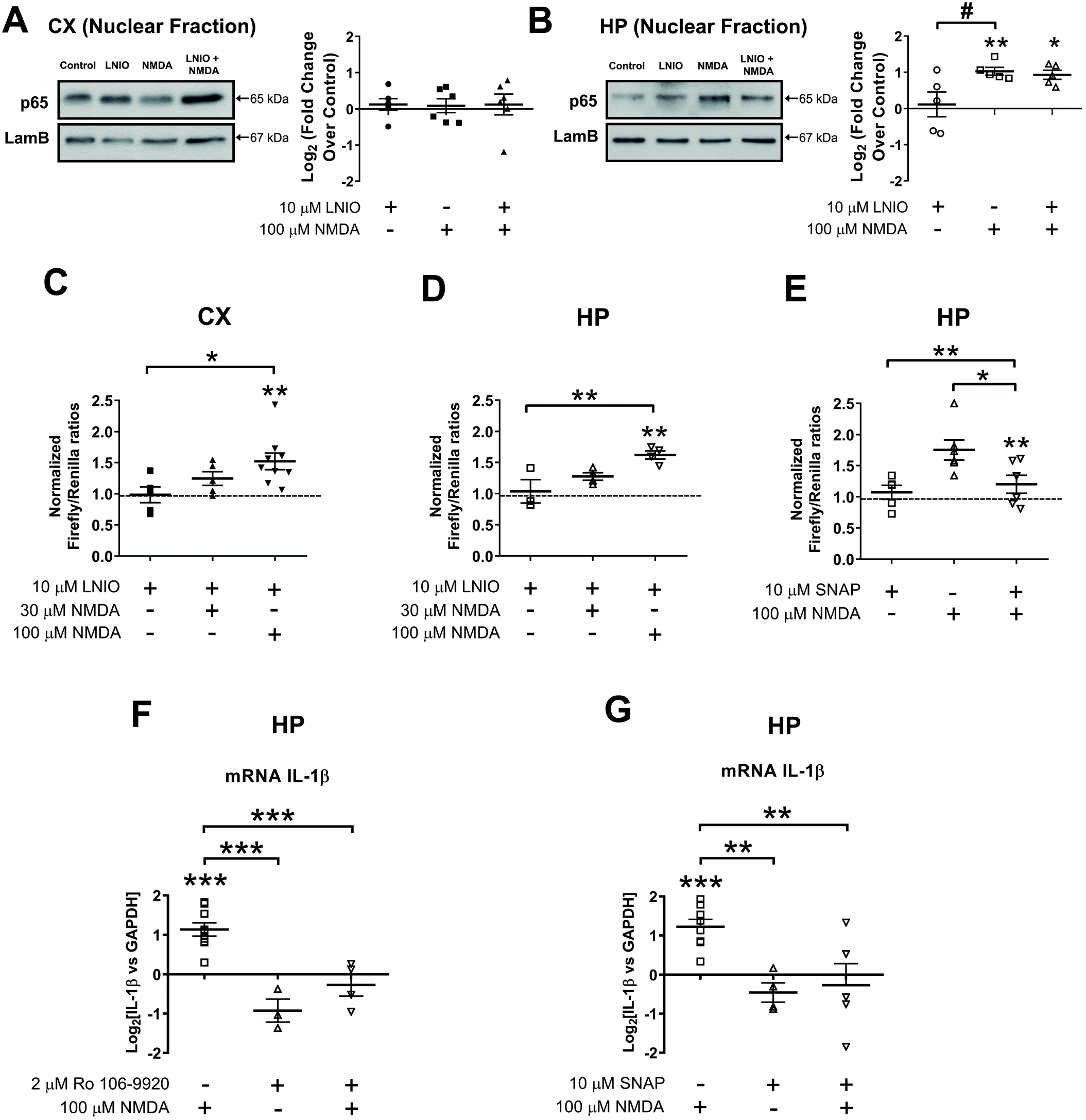
Nitric monoxide decreases transcriptional activity and gene expression but not nuclear translocation of NF-κB in response to NMDA stimulation. **A) and B)** Representative Western blots and densitometric quantifications of nuclear content of p65 in cortical (A) and hippocampal (B) cultures stimulates with NMDA (100 μM) in presence or absence of NO inhibitor LNIO (N5-(1-Iminoethyl)-L-ornithine). For each Western blot, equal quantities of protein were loaded and Lamin B1 (LamB) was used as loading control for nuclear fraction. All results were obtained in n=5 independent experiments. * p<0.5; ** p<0.1 by two-way ANOVA followed by Bonferroni post-test. **C) and D)** Relative luciferase activity in cortical (C) and hippocampal (D) neurons after 30 μM and 100 μM NMDA stimulation in presence and absence of LNIO. E) Relative luciferase activity in hippocampal neurons after NMDA 100 μM stimulation in presence and absence of NO donor SNAP at 10 μM (S-nitroso-N-acetylpenicillamine). All results were obtained in n=6 to 10 independent experiments. Statistical significance was assessed by One-way ANOVA followed by Bonferroni post-test. * p<0.5; ** p<0.1; *** p<0.001. F and G) IL-1B mRNA measured by quantitative PCR in hippocampal cultures 2 hours after NMDA 100 μM stimulation in presence or absence of 2 μM Ro 106-9920 or 10 μM SNAP. Bar graph showing the mean ± SEM fold change normalized against GAPDH as reference. Data obtained from 4 to 8 independent hippocampal cell culture experiments. Statistical significance was assessed by One-way ANOVA followed by Bonferroni post-test. **p<0.01; ***p<0.001.

To further investigate whether NOS inhibition affected the transcriptional activity of NF-κB, we used the luciferase reporter system (Figures 5 C-D). In cortical cultures, the presence of 10 μM LNIO led to increased transcriptional activity after 100 μM NMDA stimulation. Similar effects were observed in hippocampal cultures. This suggests that NOS-dependent NO synthesis leads to NF-κB inhibition. Consistently, the NO donor SNAP (10 μM) had an inhibitory effect on NF-κB activity after 100 μM NMDA (Figure 5E)^16,30^.

Finally, by measuring mRNA levels of known NF-κB downstream pro- or anti-apoptotic genes (BAX, Caspase 11, Bcl_2_) using qPCR, we investigated whether NF-κB activation after 100 μM NMDA in hippocampal neurons was associated with enhanced transcription. Surprisingly, we did not detect any changes in the mRNA levels of these genes (not shown), while changes were observed in the mRNA levels of the pro-inflammatory cytokine IL-1β. In time course experiments, we could detect that IL-1β increased after 2 hours of stimulation with 100 μM NMDA (Supplemental Figure S4), and this was inhibited in the presence of the NF-κB inhibitor Ro 106-9920 (2 μM) (Figure 5F). In a different set of experiments, it was observed that the NO donor SNAP (10 μM) also inhibited the increased transcription of IL-1β after 100 μM NMDA (Figure 5G). These results suggest that NF-κB activation in hippocampal neurons induces the transcription of the pro-inflammatory cytokine IL-1β, while this can be prevented using a NO donor to promote the inhibitory S-nitrosylation of NF-κB. Alternatively, other regulatory proteins of the NF-κB pathway could also be NO targets. In order to assess whether S-nitrosylation can be considered a more general mechanism regulating the outcome of excitotoxic stimuli, we analyzed the S-nitrosylated proteome in cortical and hippocampal cultures after NMDA.

### Detection of S-nitrosylated proteins by mass spectrometry

Hippocampal and cortical cultures were incubated in the presence or absence of 30 μM NMDA to pull down S-nitrosylated proteins using the biotin switch assay (Figure 6). Interestingly, we found that, in hippocampal neurons, a lower number of proteins were detected (178 proteins in hippocampal neurons *versus* 360 proteins cortical neurons) (Figure 6A and Supplemental Table 2). To exclude technical issues resulting in the detection of lower number of proteins in hippocampal cultures, we carefully ascertained that equal quantities of inputs were used (*i.e*., Supplemental Figure S1). These results suggest that protein S-nitrosylation levels are elevated in cortical neurons, both under control and excitotoxicity conditions, compared to hippocampal cultures. The respective Venn diagrams (Figure 6C) revealed that in cortical cultures, 41 and 64 proteins, respectively, were identified exclusively in control or 30 μM NMDA-stimulated cortical cultures, while in hippocampal neurons (Figure 6D), 8 and 40 exclusive proteins were found. After 30 μM NMDA exposure, 226 proteins were restricted to cortical and 77 proteins to hippocampal cultures (Figure 6 B). To find out which biological processes were selectively affected by NMDA in both culture types, a meta-analysis using the protein lists obtained after NMDA stimulation revealed that different biological processes were affected in each case (Figure 6E). Interestingly, in cortical cells, the S-nitrosylation (and consequent inhibition) of the proteasome subunits may contribute to decreased proteasomal degradation of the NF-κB inhibitor IκBα, thus providing an additional level of NF-κB inhibition in cortical excitotoxicity^31,32^. On the other hand, in hippocampal neurons, a functional cluster involved in actin filament capping or brain development stands out. In neurons, the actin cytoskeleton plays a major role in membrane remodeling, organelle trafficking and excitotoxicity^19,33^. The role of S-nitrosylation of actin cytoskeleton associated regulatory or motor proteins has not yet been assessed in neurons, although in cardiomyocytes, their S-nitrosylation leads to inhibition, *i.e*., lower calcium sensitivity and decreased muscle contraction^34,35,36^.

**Figure 6.**
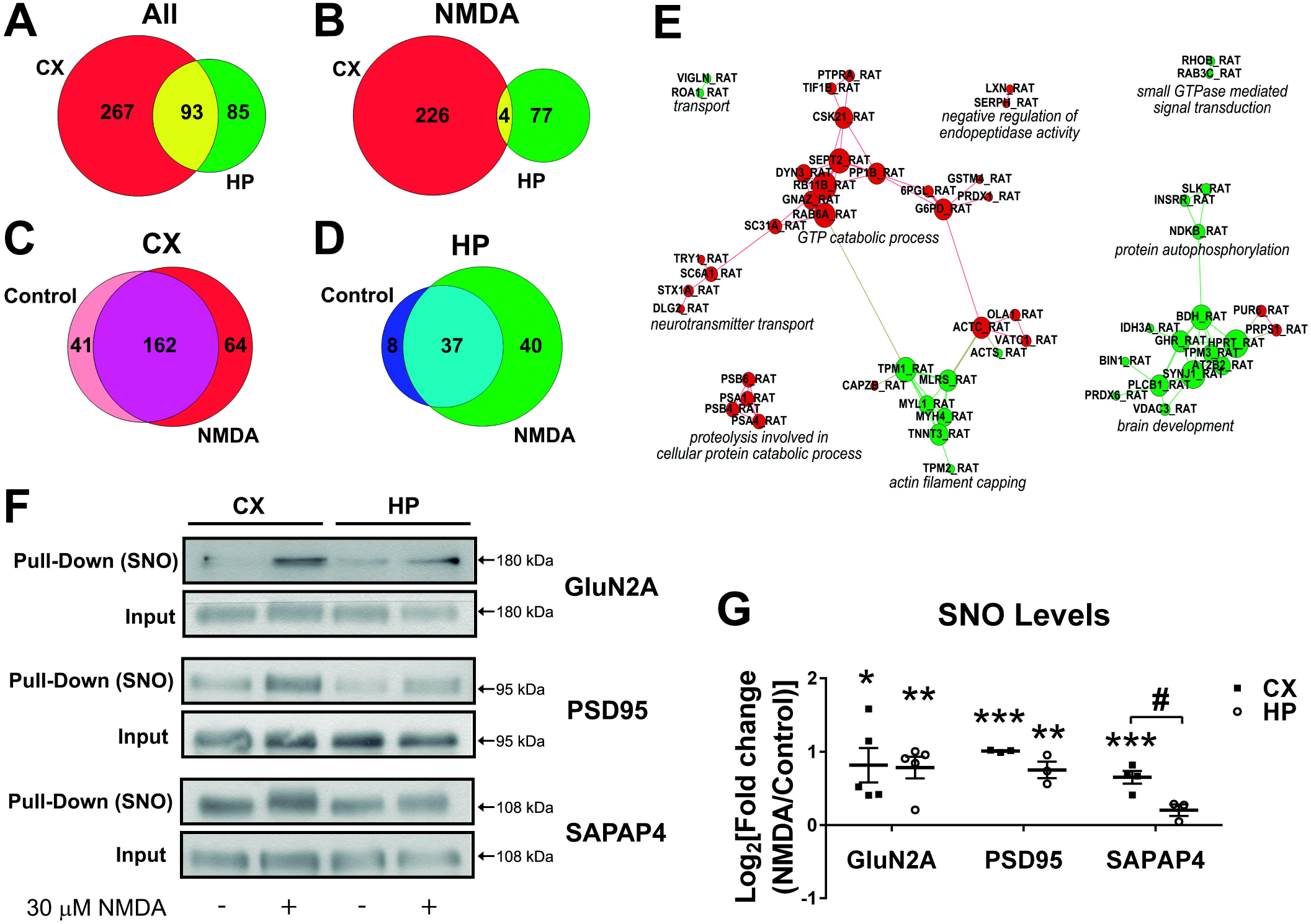
Identification of S-nitrosylated proteins after NMDA stimulation by nanoLC-MS/MS. S-nitrosylated proteins in hippocampal and cortical cultures were identified by mass spectrometry after 30 μM NMDA (n=6 except for hippocampal neurons incubated with NMDA (n=5)). A remarkable larger number of S-nitrosylated proteins were detected in cortical than in hippocampal neurons. **A)** Venn Diagrams showing the distribution of proteins in cortical vs. hippocampal cultures (using the sum of identified proteins in both, control and NMDA stimulated cultures). **B)** cortical vs. hippocampal cultures, using proteins identified under NMDA stimulation. **C)** Proteins identified in control vs NMDA stimulated cortical cultures **D)** Proteins identified in control vs NMDA stimulated hippocampal cultures. In C and D, only proteins were considered which were detected at least two times under each experimental condition. CX= cortical cell cultures, HP=hippocampal cell cultures. **E)** Meta-analysis of proteomic data using GeneCodis. Identified proteins exclusively detected in cortical (red) or hippocampal (green) proteomes were functionally annotated using the web-based tools GeneCodis and Gene Ontology (GO). A single enrichment analysis of biological processes was performed with each list of proteins. The obtained data were visualized by building a graph where the nodes are the proteins that are annotated with the enriched biological processes terms from Gene Ontology. The connections where made by looking at the enriched terms the Proteins where annotated with. If two proteins had the same annotation in common, a line was drawn. When two different colored nodes *i.e*., proteins are not connected they don’t share the same biological processes. To emphasize the similarities a force field embedder was used to layout the graph, depicting similar proteins closer to each other. Note that S-nitrosylation controls different cellular pathways. **F)** Validation by Western blot of S-nitrosylated proteins that were pulled down with the biotin switch method. **G)** Densitometric quantification of S-nitrosylated (-SNO) proteins in cortical (CX) and hippocampal (HP) cultures after NMDA stimulation, in n=4 independent experiments. # p<0.05; * p<0.05; ** p<0.01; *** p<0.001.

We also determined whether a difference in protein S-nitrosylation between both culture types could be detected in already well-validated NO targets. Thus, we quantified the S-nitrosylation of the NMDA receptor subunit GluN2A^37^ and the scaffolding protein PSD95^38^. Furthermore, we included the synapse associated protein SAPAP4, a scaffolding protein that had been detected by us in a previous S-nitrosyl proteome (unpublished) (Figure 6F-G). The S-nitrosylation of the synaptic proteins GluN2A and PSD95 was increased in both culture types after NMDA. Interestingly, S-nitrosylated SAPAP4 increased in cortical cultures, while no changes were observed in hippocampal cells, showing that in addition to p65, NO has different protein targets in the two cell types.

Finally, our results can be summarized in the model presented in Figure 7.

**Figure 7.**
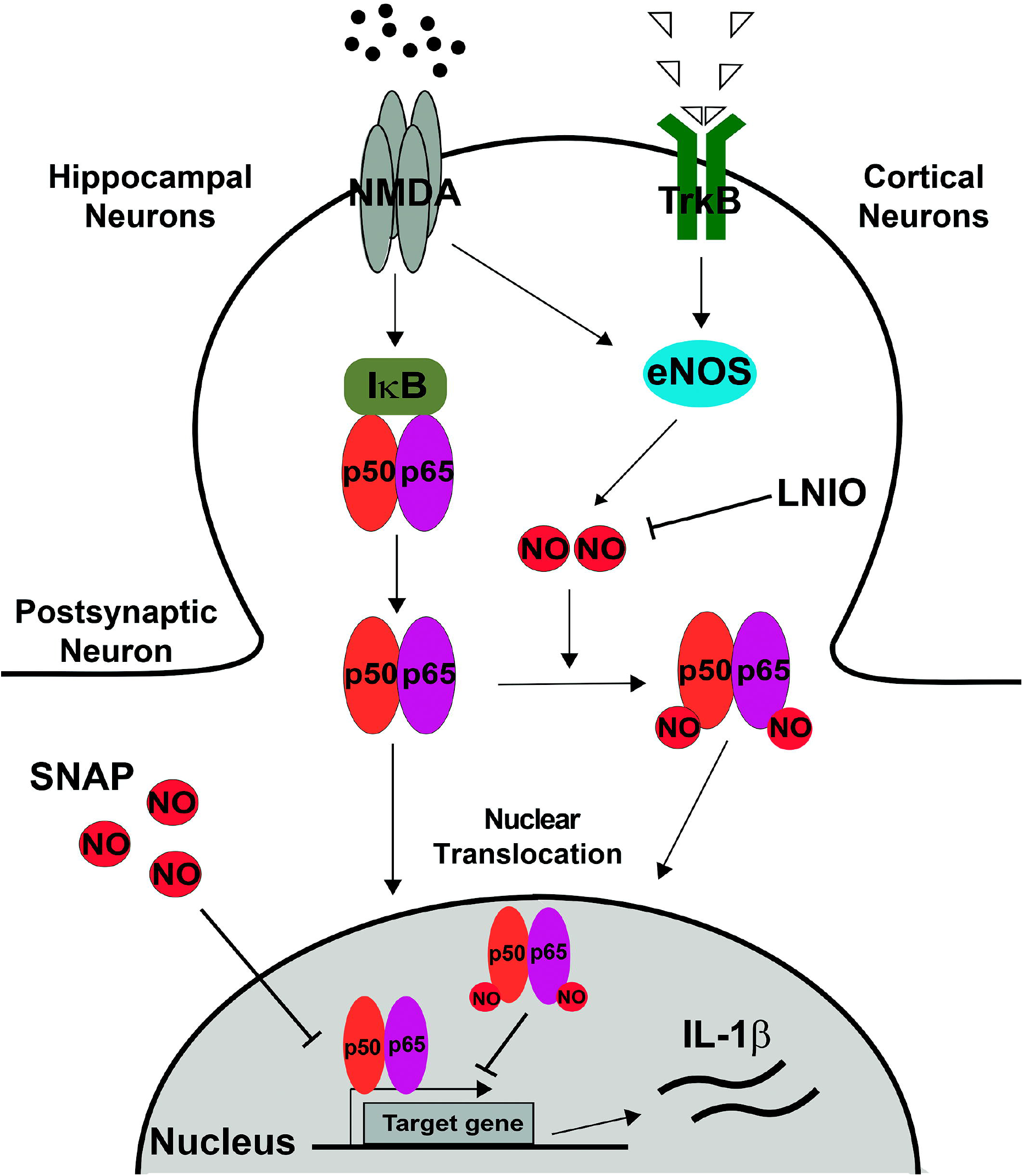
Proposed model summarizing the results. Induced/activated eNOS located at the excitatory synapse produces NO, leading beside others to NF-κB S-nitrosylation in cortical cells and inhibiting NF-κB-dependent gene expression. However, under excitotoxic conditions this eNOS-dependent negative regulation of p65 is not present in hippocampal cultures. Therefore, NMDA leads to the activation and nuclear translocation of NF-κB, resulting in a transcriptional activation that includes pro-inflammatory genes, including IL-1β. The transcriptional activity of NF-κB can be selectively induced in cortical cultures by inhibiting NOS enzymes with LNIO. Likewise, in hippocampal cultures, the transcriptional activity (including IL-1β transcription) can be inhibited by the NO donor SNAP.

## Discussion

In this work, we show that eNOS-dependent p65 S-nitrosylation after NMDA receptor overstimulation is neuroprotective. Previously, NO has been proposed as a promising therapeutic target for dealing with excitotoxic insults in the developing brain^39^. Moreover, in several preclinical models of ischemic stroke followed by reperfusion, or of traumatic brain injury, increasing eNOS-dependent NO production or the cerebral NO levels, either by using NO donors or NO inhalation, has neuroprotective effects^40^. In recent studies, NO-mediated protection has been shown in cerebellar granule neurons^41^ and in a testicular ischemia/reperfusion model^42^ while eNOS-dependent NO production protected the neurovascular unit and ameliorated neurological deficits^43,44^. Moreover, NO was neuroprotective in various animal models of Parkinson Disease, after oxygen glucose deprivation or cerebral ischemia/reperfusion injury and this effect depended on a reduction in reactive oxygen species and protein S-nitrosylation in brain mitochondria^45,46^, while in a pharmacological study, neuroprotection occurred in a PI3K/Akt dependent manner^47^. Interestingly, NO-signaling deficiency may contribute importantly to age-related cognitive impairment^48^. In turn, and in accordance with our data, brain ischemia induced a deleterious elevation of NO and NOS in the hippocampus^49^. Thus, our results add to our understanding of neuronal mechanisms that participate in NO mediated neuroprotection, which, we hope, will help in the development of novel therapeutic strategies aimed at inhibiting harmful NF-κB activity in acute and chronic neurodegenerative disorders^50^.

### NF-κB and eNOS-dependent NO production in the cerebral cortex

The neurotrophin BDNF and its tropomyosin-related kinase receptor TrkB, a signaling system associated importantly with improvement of cognitive functions in the central nervous system, is known to activate eNOS in endothelial cells^51,52^. Thus, we used BDNF to stimulate eNOS-dependent neuroprotective NO synthesis in our cell model^16^. Consistent with our results, the restitution of BDNF/TrkB signaling after a stroke enhanced neuroprotection in the cerebral cortex^53^. Moreover, further recent studies have confirmed functional implications of eNOS expression in neurons^54^. We focused on NF-κB, a known target of NO and also implicated in both neuroprotective and neurotoxic effects.

Constitutive NF-κB activity has been described in the cerebral cortex, hippocampus, amygdala, cerebellum, hypothalamus and olfactory bulbs^7,8^. In *in vivo* experiments using a transgenic mouse model in which NF-κB expression was measured by β-galactosidase activity, high constitutive expression was found in the CA1, CA2 and dentate gyrus regions of the hippocampus, while lower levels were found in the cerebral cortex^55^. This constitutive activity is beneficial for neuronal survival, as well as for learning and memory, and, thus, might favor the transcription of genes involved in these processes. However, NF-κB activation favored cell death or damage in pathophysiological models that involve NMDA receptor overactivation^56,57,58^ while it resulted neuroprotective in cortical neurons both in vitro and in vivo^21^. It is unknown how an excitotoxic insult might switch NF-κB activity to promote the expression of deleterious or pro-inflammatory proteins^59^. One possibility is that different post-translational modifications that act in concert, also known as the “bar code” for NF-κB activation, determine this switch^60^. In such a way, the interaction of S-nitrosylation with phosphorylation, which is importantly regulated under excitotoxic conditions, remains unexplored^19^.

In addition to p65, the p50 subunit of NF-κB can be S-nitrosylated at the highly conserved cysteine 62 residue, and, similarly to p65 modification, this results in the inhibition of its DNA binding capacity, contributing to NF-κB inhibition^10,11,61^. Another component of the NF-κB pathway that can be S-nitrosylated is the inhibitor of NF-κB (IκB) kinase (IKK) complex, the main kinase complex responsible for the phosphorylation of the IκB-α protein. The IKK complex is composed of the two catalytic subunits IKK-α and IKK-β and the regulatory subunit IKK-γ. The S-nitrosylation of the cysteine 179 residue of the IKK-β subunit results in the inhibition of the kinase activity of the IKK complex and consequently the lack of IκB-α protein phosphorylation, thus preventing activation of NF-κB^62^. In consequence, enhanced protein S-nitrosylation of different NF-κB pathway components converge on its inhibition. Because of the dearth of NF-κB molecules and their regulators compared to other proteins, *e.g*., those of the cytoskeleton, we failed to detect them on the mass spectrometric screens of S-nitrosylated proteins. Remarkably, even the most up-to-date and most sensitive approach to demonstrating S-nitrosylation (*i.e*. Cys-BOOST, bio-orthogonal cleavable-linker-based enrichment and switch technique), was not capable of detecting any NF-κB associated molecules so far^63^. Moreover, when separating neuronal cell nuclei to obtain enrichment of S-nitrosylated nuclear proteins and a higher chance to detect less abundant proteins, NF-κB remained hidden^64^.

### The SNO proteome after excitotoxicity

S-nitrosylation of proteins is the principal cGMP-independent mode of action of NO. The S-nitrosylation of redox-sensitive cysteins has been described in thousands of proteins that regulate a variety of biological functions^63,65^. In total, our MS-based S-nitrosylation screen identified 445 different proteins. Hierarchical gene ontology (GO)-based clustering of those proteins (Supplemental Table 3) revealed a strong participation in metabolic processes, including glycolysis, tricarboxylic acid cycle, 2-oxoglutarate process, ATP biosynthetic process and carbohydrate metabolic process. This ranking was followed by increased S-nitrosylation of mitochondrial proteins modulating their function, including negative effects on the electron transport chain, alteration in the mitochondrial permeability transition pore and enhanced mitochondrial fragmentation and autophagy^66^. However, proteins participating in neuron projection development and brain development as well as in synapse associated processes, with roles in synaptic transmission, neurotransmitter transport and ionotrophic glutamate receptor signaling, are within the top 35 of this list. This indicates that, besides metabolic processes, even basic neuronal mechanisms are regulated by S-nitrosylation. The current view is that under conditions of normal NO production, S-nitrosylation regulates the activity of many normal proteins; however, increased levels of NO, as experimentally induced by lasting NMDA stimulation, led to aberrant S-nitrosylation, thus contributing to the pathogenesis of neurodegenerative disorders^67^. Remarkably, in this context, we found increases in the GO terms “protein phosphorylation” and “protein autophosphorylation” (Supplemental Table 3) after NMDA. S-nitrosylated proteins belonging to these terms include important serine kinases, including CaMK2d, GSK3β, Akt1 and MAPKinases, but also tyrosine kinases like Fyn and Src. This result indicates that regulation of kinase activity by S-nitrosylation might contribute to the NMDA-induced phosphoproteome^19^ and in the case of NF-κB, this would contribute to the generation of the “bar code” specifying its transcriptional targets. For example, S-nitrosylation of Src overrides an inhibitory phosphorylation motif leading to a phosphorylation independent activation of this kinase^68,69^. Moreover, S-nitrosylation of CaMKII, a central neuronal kinase implicated synaptic plasticity, can induce its Ca^2+^independent activation^70^, while the opposite effect, was also described^71^. But it is beyond doubt that S-nitrosylation can strongly modulate the activity of key kinases in neurons that, in turn, are known NF-κB regulators^8,72^.

Our results show that sustained NMDA receptor activation results in a substantially modified S-nitrosylation proteome in neurons. In them, protein clusters that regulate the NF-κB pathway were found, *e.g*., S-nitrosylation of proteasomal proteins causes its inhibition and, therefore, decreased degradation of IκB should be expected, thus contributing to NF-κB inhibition^31,32^. The work presented here encourages therapeutic strategies directed to favor homeostatic adaptation associated to NMDA receptor overstimulation, an idea that is supported by the positive effects of NF-κB inhibition in aging in increasing health and lifespan^50^.

## Methods

### Material

Chemical reagents were purchased from Sigma (St. Louis, MO, USA), unless otherwise stated. Neurobasal medium (Cat. N°: 21103-049), B27 (Cat. N° 17504-044), MEM (Minimum Essential Medium Cell Culture) (Cat. N° 11900-024), FBS (Fetal Bovine Serum) and Equine Serum (Cat. N° 16050-122) were obtained from Gibco-Invitrogen (San Diego, CA, USA). Penicillin-Streptomycin was obtained from Biological Industries (Cromwell, CT, USA). N-Methyl-D-aspartate (NMDA) (Cat. N° 0114), 6-Cyano-7-nitroquinoxaline-2,3-dione (CNQX) (Cat. N° 0190) and N5-1(1-Iminoethyl)-L-ornithine dihydrochloride (LNIO) (Cat. N° 0546) were obtained from Tocris Bioscience (Bristol, UK). 2-amino-5-phosphonovalerate (APV) (Cat. N° A-169) was obtained from RBI (Natick, MA, USA). Recombinant Escherichia coli-derived BDNF was obtained from Alomone Labs (Jerusalem, Israel). Ro 106-9920 (6-(Phenylsulfinyl) tetrazolo[1,5-b] pyridazine) (Cat. N° 1778), Nimodipine (Cat. N° 482200), S-nitroso-N-acetylpenicillamine (SNAP) (Cat. N° 487910) and 3-amino,4-aminomethyl-2’,7’-difluorofluorescein (DAF-FM) (Cat. N° 251515) were obtained from Calbiochem (San Diego, CA, USA). EZ-link HPDP-Biotin (Cat. N° 21341) and Streptavidin Agarose (Cat. N° 20347) were obtained from Thermo Scientific, (Waltham, MA, USA). Trypsin Gold was obtained from Promega (Cat. N° V5280) (Madison, WI, USA).

### Antibodies

*Primary antibodies:* Anti-p65 (Cat. N° ab16502), Anti-IκB alpha (Cat. N° ab32518), Anti-Laminin-B1 (Cat. N° 8982), Anti-Tubulin Alpha 1A (Cat N° ab7291) and Anti-GAPDH (Cat. N° ab8245) were from Abcam (Cambridge, UK). Anti-phospho-p65 was obtained from Cell signaling (Cat. N° 3033) (Danvers, MA, USA), Anti-MAP2A/2B was obtained from Millipore (Cat. N° MAB378) (Burlington, MA, USA), Anti-GFAP was obtained from US Biological (Cat. N° G2032-28B-PE) (Swampscott, MA, USA), Anti-βIII tubulin was obtained from Promega (Cat. N° G712A) (Madison, WI, USA), Anti-GluN2A was obtained from Alomone Labs (Cat. N° AGC-002) (Jerusalem, Israel), Anti-SAPAP4 was obtained from Santa Cruz Biotechnology (Cat. N° sc-86851) (Dallas, TX, USA), Anti-Biotin was obtained from Bethyl laboratories (Cat. N° A150-111A) (Montgomery, TX, USA) and Anti-PSD-95 was obtained from BD transduction Laboratories (Cat. N° 610495) (San Jose, CA, USA). *Secondary Antibodies:* HRP Goat anti Rabbit IgG (Cat. N° 926-80011) and HRP Goat anti-Mouse IgG (Cat. N° 926-80010) were from LI-COR Biosciences (Lincoln, NE, USA), Alexa Fluor^®^ 555 goat anti rabbit IgG (Cat. N° A21429) was obtained from Life Technologies (Carlsbad, CA, USA), Alexa Fluor® 488 Goat Anti-Mouse IgG (Cat. N° A21202) was obtained from Invitrogen Corporation (Carlsbad, CA, USA).

### Neuronal cultures

Primary cultures of cortical (CX) and hippocampal (HP) neurons were obtained from day-18 Sprague-Dawley rat embryos, as described^16^. Procedures involving animals and their care were approved by the Universidad de los Andes Bioethical Committee and performed in accordance to the ARRIVE Guidelines. Neurons were cultured in the absence of Cytosine arabinoside (AraC) and contained about 30% of astrocytes^17^. The excitotoxic stimulation was induced by addition of 30 to 100 μM NMDA and 10 μM glycine for 60 minutes. When indicated, the NMDA stimulus was applied after a 15 min preincubation with 10 μM LNIO, 2 μM Ro 106-9920 or 10 μM SNAP.

### Cell fractionation

Cell fractionation was performed immediately after the excitotoxic stimulation (NMDA + glycine for 1 hour). Cells were harvested in buffer A (0.6% NP40 v/v; in mM: 150 NaCl; 10 HEPES pH 7.9; 1 EDTA) and homogenated in Teflon-glass homogenizer, vortexed for 30 seconds and incubated on ice for 10 minutes. This procedure was repeated 3 times. The suspension was centrifuged at 17,000 g by 5 minutes to obtain the cytoplasmic fraction. The pellet was washed with buffer B (in mM: 150 NaCl; 10 HEPES pH 7.4; 1 EDTA) and centrifuged at 17,000 g by 1 minute at 4°C, resuspended in buffer C (25% v/v glycerol; in mM 20 HEPES pH 7.4; 400 NaCl; 1.2 MgCl_2_; 0.2 EDTA), vortexed for 30 seconds and incubated on ice for 10 minutes (5 times) to finally centrifuge at 17,000 g for 20 minutes to obtain the nuclear fraction.

### Cell viability

The percentage of surviving neurons was assessed 24 h after the NMDA challenge using the trypan blue exclusion test, in 24 well plates containing 10,000 cells. Neurons were exposed to 0.05% (v/v) trypan blue in PBS for 5 minutes. The cells were immediately examined under a phase-contrast microscope, images of ten random fields were recorded to quantify the numbers of living neurons (which exclude trypan blue) and dead (stained) neurons.

### Immunocytochemistry

Neuronal cultures of 14 to 15 DIV were fixed immediately after the excitotoxic insult with 4% paraformaldehyde in PBS containing 4% of sucrose for 10 minutes and washed with PBS. After fixation the cells were permeabilized with 0.2% Triton X-100 for 5 minutes and washed with PBS containing 25 mM glycine. Cells were incubated with blocking solution (10% BSA in PBS) for 1 h followed by overnight incubation with primary antibody: anti-p65 (1:300), anti-MAP2A/2B (1:1000) and anti-GFAP (1:1000), all diluted in the same blocking solution at 4°C. After incubation with primary antibody, cells were washed with PBS, blocked for 30 minutes with 10% BSA and incubated for one hour at room temperature with the corresponding secondary antibody diluted 1:1000 in blocking solution and finally incubated with DAPI for 5 minutes for nuclear staining. The fluorescence images were obtained using ECLIPSE TE2000U Microscope with NIS-Element imaging software from Nikon Instrument Inc (Minato, Tokio, Japan), and analyzed using Photoshop CS6 software. In order to assess the nuclear translocation of NF-κB by epifluorescence microscopy, 50 cells per condition (control or NMDA) were analyzed in which the nuclear (*i.e*., DAPI stained) zone was selected and the intensity of p65 was quantified in that area by an experimenter blind to the experimental conditions. Finally, the decodification of the data allowed the comparison of fluorescence intensity of p65 in control and NMDA stimulated cultures.

### Nitric oxide production

Neuronal cultures were loaded for 1 h at 37°C with 10 μM 4-amino-5-methylamino-2’,7’-difluorofluorescein (DAF-FM) plus 0.015% pluronic acid in recording solution (in mM: 116 NaCl, 5.4 KCl, 0.9 NaH2PO4, 1.8 CaCl_2_, 0.9 MgCl_2_, 20 HEPES, 10 glucose and 0.1 L-arginine, pH 7.4). Cells were washed 4 times and placed in recording solution. Fluorescence (excitation at 495 nm; emission at 510 nm) were acquired for 500 ms every 5 minutes to minimize the photobleaching of DAF-FM^73^. Signals were averaged over regions of interest of somas (excluding the nuclei) and relative intracellular NO levels were calculated from emission at 510 nm. Because there was a linear decay of fluorescence due to photobleaching, the negative slope was determined for each experiment before the addition of the stimulus (BDNF), and the experimental slope was corrected for this^16^. At the end of the experiment, the external NO donor S-Nitroso-N-acetyl-DL-penicillamine (SNAP, 10 μM) was applied to check that NO-sensitive dye was still available. Experiments in which SNAP did not increase fluorescence were discarded. Fluorescence was measured using an Eclipse E400 epifluorescence microscope with a FluorX40 water immersion objective (Nikon Corporation, Melville, NY, USA) equipped with a Sutter Lambda 10-2 optical filter changer. Emitted fluorescence was registered with a cooled charge-coupled device video camera (Retiga 2000R Fast 1394, QImaging, Surrey, BC, Canada) and data obtained were processed using imaging software (IPLab 4.0, Scanalytics, Buckinghamshire, UK).

### High resolution proteome analysis and label free quantitation

The proteins pulled down in the biotin switch assay were boiled in denaturizing SDS-sample buffer and subjected to SDS-PAGE (n=6 biological replicates for each experimental condition except for hippocampal neurons incubated with NMDA (n=5)). SDS-gels (3% stacking gel, 12% separation gel) were run in a Mini PROTEAN® System (BioRad) at 100 V for 10 min and 200 V till end of the separation. Each lane was divided in eight fractions for in-gel-digestion and further analysis. In-gel digest was performed in an adapted manner according to Shevchenko^74^. LC-MS/MS analyses of the generated peptides were performed on a hybrid dual-pressure linear ion trap/orbitrap mass spectrometer (LTQ Orbitrap Velos Pro, Thermo Scientific, San Jose, CA) equipped with an EASY-nLC Ultra HPLC (Thermo Scientific, San Jose, CA). Peptide samples were dissolved in 10 μl 2% ACN/0.1% trifluoric acid (TFA) and fractionated on a 75 μm I.D., 25 cm PepMap C18-column, packed with 2 μm resin (Dionex, Germany). Separation was achieved through applying a gradient from 2% ACN to 35% ACN in 0.1% FA over a 150 min gradient at a flow rate of 300 nl/min. The LTQ Orbitrap Velos Pro MS has exclusively used CID-fragmentation when acquiring MS/MS spectra consisted of an Orbitrap full MS scan followed by up to 15 LTQ MS/MS experiments (TOP15) on the most abundant ions detected in the full MS scan. Essential MS settings were as follows: full MS (FTMS; resolution 60.000; m/z range 400-2000); MS/MS (Linear Trap; minimum signal threshold 500; isolation width 2 Da; dynamic exclusion time setting 30 s; singly-charged ions were excluded from selection). Normalized collision energy was set to 35%, and activation time to 10 ms. Raw data processing and protein identification of the high resolution Orbitrap data sets was performed by PEAKS software suite (Bioinformatics Solutions, Inc., Canada). False discovery rate (FDR) was set to < 1%.

### Western blotting

Twenty micrograms of protein of each sample, dissolved at 1 mg/ml in loading buffer, were separated by sodium dodecyl sulfate-polyacrylamide electrophoresis (SDS-PAGE) on 10% gels under fully reducing conditions and transferred onto nitrocellulose membranes. Membranes were incubated overnight at 4°C with primary antibodies followed by incubation at room temperature with secondary antibody conjugated with horseradish peroxidase for 60 min. Immunoreactivity was visualized using the ECL detection system. Densitometric quantification was performed using the image processing program ImageJ (National Institute of Health, USA). Data were expressed as fold change from homogenate for at least 4 independent preparations and mean ± SEM for each fraction was calculated.

### Quantitative PCR

Total RNA from primary hippocampal cultures was extracted using TRizol reagent from Life technologies (Carlsbad, CA, USA), 1 μg of RNA was reverse transcribed into cDNA using MultiScribe reverse transcriptase from ThermoFisher (Waltham, MA, USA) according to the manufacturer’s protocol. Quantitative polymerase chain reaction (qPCR) reaction was carried out using the Brilliant III Ultra-Fast QPCR Master Mix in the Stratagene Mx3000P system (Agilent Technologies, Santa Clara, CA, USA). The thermal cycling protocol was: pre-incubation, 95°C, 10 min; amplification, 40 cycles of (95°C, 20 s; 60°C, 20 s; 72°C, 20 s); melting curve, 1 cycle of (95°C, 1 min; 55°C, 30 s; 95°C, 30 s). qPCR was performed using triplicates. Primers used were: rat IL-1β, forward primer 5’ TCAGGAAGGCAGTGTCACTCATTG 3’ and reverse primer 5’ ACACACTAGCAGGTCGTCATCATC 3’. The results were normalized against rat mRNA of GAPDH, Forward primer 5’ TTCACCACCATGGAGAAGGC 3’ and reverse primer 5’ GGCATGGACTGTGGTCATGA 3’. The gene expression was represented by the value of ΔCt (Sample Problem Ct – Reference Gene Ct). The relative expression is expressed as fold change over control using the 2-^ΔΔCt^ expressed on base 2 logarithmic scale.

### Knockdown of eNOS

Short hairpin against eNOS (sh-eNOS) was synthesized in integrated DNA technologies (IDT) (Neward, NJ, USA), aligned and expressed in the lentiviral vector pLL3.7-mRuby2, downstream of the U6 promoter and between HpaI and XhoI sites. The sh-eNOS sequence was: 5’-GTGTGAAGGCGACTATCCTGTATGGCTCT-3’. The scrambled RNA (sh-Luc) sequence was: 5’-TTCTCCGAACGTGTCACGT-3’. Correct insertions of the shRNA cassettes were confirmed by restriction mapping and direct DNA sequencing. Lentiviral production was done using lipofectamine 2000 reagent, Promega (Cat. N° 11668-019) (Madison, WI, USA). Briefly, we co-transfected the sh-eNOS or sh-Luc plasmids with the packaging vector Δ8.91 and the envelope vector VSV-g into HEK293T cells in free serum DMEM. 5 hours after transfection the medium was replaced for DMEM containing 10% FBS and the next day the medium was replaced by Neurobasal supplemented with B27. The resulting supernatant contained the lentiviruses (Naldini et al., 1996; Dull et al., 1998).

### Magnetofection of primary neurons

Neuronal cultures of 7 DIV were transfected using magnetic nanoparticles (NeuroMag, Oz Biosciences). Briefly, plasmid DNA of Firefly and Renilla Luciferase were incubated with NeuroMag Transfection Reagent (in a relationship of 2 μl per 1 μg of DNA) in Neurobasal medium, added to the cultures to incubate for 15 minutes at 37°C on the magnetic plate.

### Dual luciferase assay

Transfected neuronal cultures with the NF-κB reporter Firefly Luciferase plasmid (Cat. N° E1980, Promega, Madison, WI, USA), were stimulated with NMDA/glycine for 60 minutes, in the presence or absence of the NO inhibitor N5-(1-Iminoethyl)-L-ornithine (LNIO). After stimulation, the cells were returned to fresh Neurobasal/B27 medium containing 10 μM CNQX, 2 μM nimodipine and 10 μM APV (to block a-amino-3-hydroxy-5-methylisoxazole-4-propionate (AMPA) receptors, Ca^2+^ channels and NMDA receptors, respectively) during 4 hours to perform the Dual-Luciferase Reporter Assay, according to the manufacturer’s protocol and carried out in FLx800 Luminometer, Biotek instrument (Winooski, VT, USA). The data were expressed as the ratio of Firefly to Renilla Luciferase activity.

### Biotin switch method

The protocol of Forrester et al. was applied with minor modifications (Supplemental Figure S1A-C)^25^. The complete procedure was performed in the dark. Neuronal cultures were homogenized in HENS buffer (in mM: 250 HEPES, 1 EDTA, 1 neocuproine, 0.1 % SDS y and protease inhibitors, pH 7.4) plus 100 mM iodoacetamide (IA). Briefly, 1 mg of starting material was blocked with 100 mM of IA in HENS buffer in a final volume of 2 ml in a rotating wheel for 1 h at room temperature, then proteins where precipitated with 3 volumes of acetone at −20°C and centrifuged at 3000 g for 10 minutes to discard the supernatant (this step was repeated two times). The blocking procedure was repeated once more. After careful resuspension, the labeling reaction was performed in the dark using 300 μl of HENS buffer containing final concentrations of 33 mM sodium ascorbate and 1 mM N-[6-(biotinamido)hexyl]-3’-(2’-pyridyldithio)-propionamide (Biotin-HPDP) (Pierce Biotechnology) biotin–HPDP. This ascorbate concentration to reduce-SNO residues falls within the wide range of concentrations suggested in the literature for ascorbate-based methods for SNO protein enrichment (i.e. from 10 to 200 mM)^34,38,75^. Then, biotinylated proteins were pulled down overnight with 200 μl of streptavidin-agarose beads in a final volume of 1 ml at 4 °C. Elution was performed with SDS gel electrophoresis loading buffer.

### Statistical Analysis

Average values are expressed as means ± SEM. Statistical significance of results was assessed using two-tailed Student’s t-test or one-way ANOVA followed by Bonferroni post-tests, as indicated. All statistic data are summarized in Supplemental Table 1.

## Acknowledgements

We thank Albert M Galaburda (Department of Neurology, Beth Israel Deaconess Medical Center and Harvard Medical School, Boston, MA) for critically reviewing the manuscript. This work was supported by Regular Fondecyt Project 1140108 to UW (Conicyt, Chile) and Regular Fondecyt Project 1200693 (ANID, Chile) to UW. Centrifugation steps were performed thanks to Project F ondequip EQM40131.

## Author Contribution

UW and TK designed the experiments and wrote the manuscript, AC prepared the final version of all figures and of the manuscript. FB, CL and KHS revised carefully the manuscript. MV designed molecular tools and supervised experiments (eNOS knockdown and dual luciferase assay). FG initiated biotin switch assay. The experimental work was done by: Figure 1, KC generated data of panels A-D, AC generated E-F; Figure 2, generated by KC; Figure 3, generated by BM; Figure 4, generated by AC; Figure 5, generated by AC and KC; Figure 6, BM did the biotin switch and generated the data of A to D, F, G; TK supervised the mass spectrometry; AE did the bio-informatic analysis.

