## Supplemental Figures and Tables for "eNOS-dependent S-nitrosylation of the NF-κB subunit p65 has neuroprotective effects"

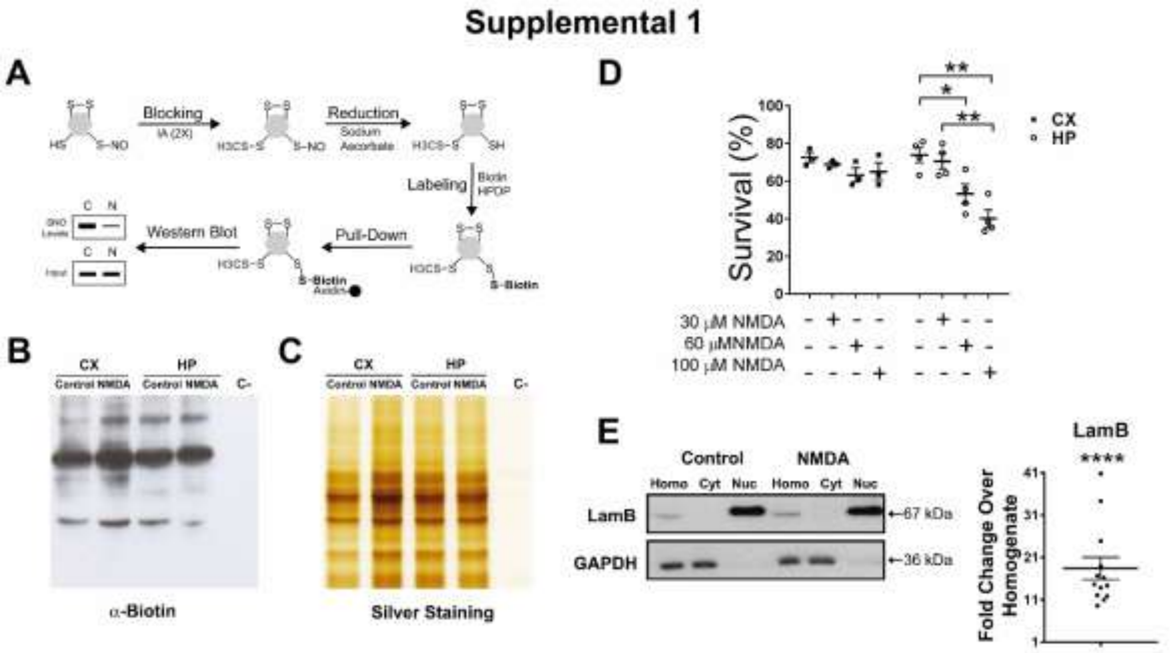


**Supplemental Figure S1. Methodological aspects: A to C: Biotin switch assay in neuronal cell cultures, D: NMDA doses and cell death in cell cultures, E: obtention of nuclear fractions.**

**A) Biotin switch assay.** Free –SH groups were blocked in 1 mg of starting material 2 times with 100 mM iodoacetamide for one hour at room temperature, followed by reduction of S-nitrosylated cysteines with 100 mM sodium ascorbate and subsequent labeling them with 300 µM biotin HPDP for one hour. Biotinylated proteins were considered as formerly S-nitrosylated ones and pulled down by streptavidin-beads. The figure was based on Forrester et al. [17]. In the negative control (C-), sodium ascorbate was omitted and thus, finally no biotinylated proteins are expected. **B) Detection of the biotinylated proteins** by Western blot using an anti-biotin antibody in cortical and hippocampal cultures, showing correct -SH group blocking and biotin labeling. **C) Silver staining of biotinylated**

**proteins** captured by streptavidin beads and separated in a 10 % SDS–PAGE. **D) Dose response curve for NMDA.** Cell viability was assessed at different NMDA concentrations, applied for one hour to measure cell death with the Trypan exclusion test 24 hours later. Hippocampal (HP) or cortical (CX) cultures were studied. 30 µM NMDA stimulation induces no significant cell death, analyzed in n= 3 to 4 independent experiment. Statistical significance was assessed by One-way ANOVA followed by Bonferroni post-test. $^{**}p<0.01$; $^{*}p<0.05$. **E) Nuclear fractionation.** Approval of sufficient purification of prepared nuclear fractions by the nuclear marker Laminin B1. Representative Western blot of total levels of Lamin B1 and GAPDH (as cytoplasmic marker) in subcellular fractions: homogenate, and the cytoplasmic and nuclear fractions of control and 30 µM NMDA stimulated neuronal cultures. An enrichment of Lamin B1 and absence of GAPDH in the nuclear fraction is observed in each case. The densitometric quantification (fold change in the nuclear fraction over homogenate) of Lamin B1 is shown (n=13 biological replicates). **** $p<0.0001$ by two-tailed t-test. Homo=homogenate; Cyt= cytoplasmic fraction; Nuc= nuclear fraction.

#### Supplemental 2

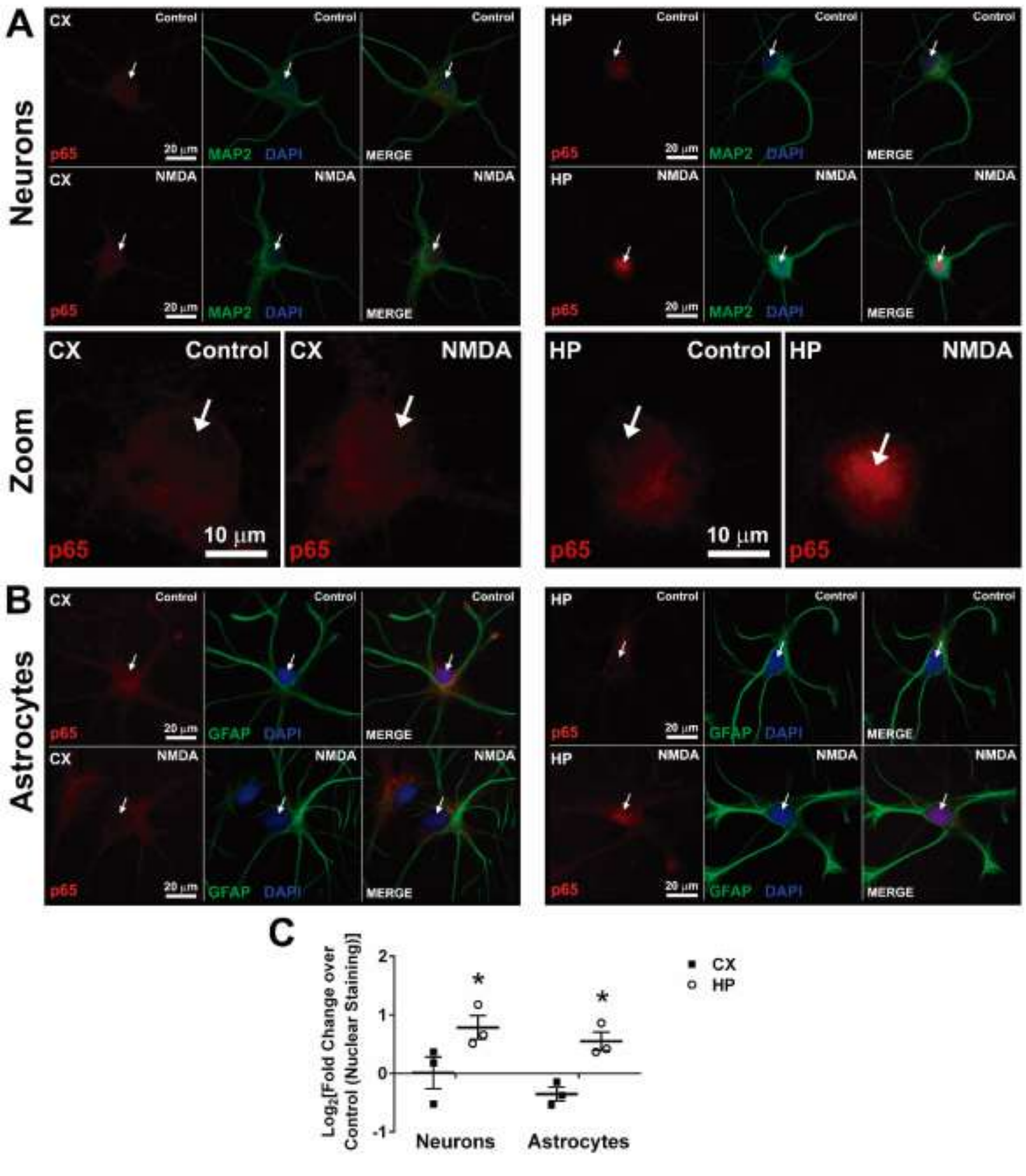

**Supplemental Figure S2.** 30 μM **NMDA induces the nuclear translocation of p65 (NF-κB) in hippocampal but not in cortical cultures.** We detected p65 in DAPI-stained nuclei of neurons (labelled with an antibody against microtubule associated protein 2, MAP2) or astrocytes (labelled with an antibody against glial fibrillary associated protein, GFAP). Neurons (top panel) and Astrocytes (Bottom panel) of cortical (left) and hippocampal (right) cultures stimulated with 30 μM NMDA for 60 minutes, were stained with antibodies against p65 (red), MAP2 (neurons, green) or GFAP (astrocytes, green). Cell nuclei were stained with DAPI (blue). The fluorescence intensity in the ROI (*i.e.*, the DAPI-positive region) was quantified. **A)** Representative images of a cortical (left) and hippocampal (right) neuronal cultures stained with p65, MAP2 and DAPI at two magnifications (calibration bars of 10 μm and 20 μM, respectively). Arrows indicate cell nuclei recognized by DAPI staining. **B)** Representative images of a cortical (left) and hippocampal (right) astrocyte cultures stained with p65, GFAP and DAPI. **C)** Relative changes of p65 fluorescence intensity in the nuclei of neurons and astrocytes of cortical (CX) and hippocampal (HP) cultures when comparing the stimulated with the control (non-stimulated) condition. Results obtained in n=4 independent experiments. * $p<0.05$ by two-tailed t-test.

### Supplemental 3

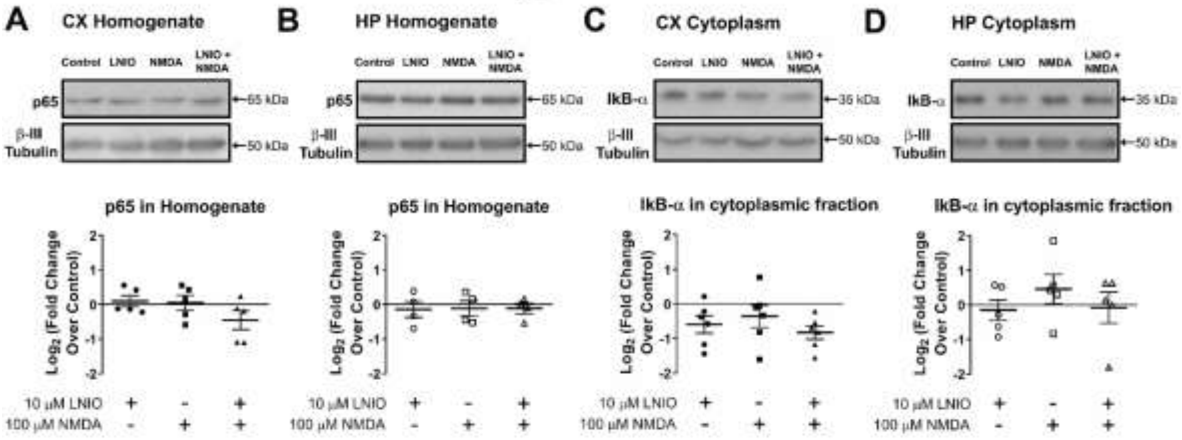


**Supplemental Figure S3. The expression levels of NF-κB subunit p65 in homogenates and of IkB-α in cytoplasmic fractions remain constant in all conditions. A) and B)** Representative Western blots and densitometric quantification of cytoplasmic content of IkB-α in cortical (A) and hippocampal (B) cultures stimulates with NMDA 100 µM in presence or absence of NO inhibitor LNIO (N5-(1-Iminoethyl)-L-ornithine). **C) and D)** Representative Wester blots and densitometric quantification of total content of p65 in cortical (C) and hippocampal (D) cultures stimulates with NMDA 100 µM in presence or absence of NO inhibitor LNIO. For each Western blot, equal quantities of protein were loaded, and β-III tubulin was used as loading control for cytoplasmic fraction and homogenate.

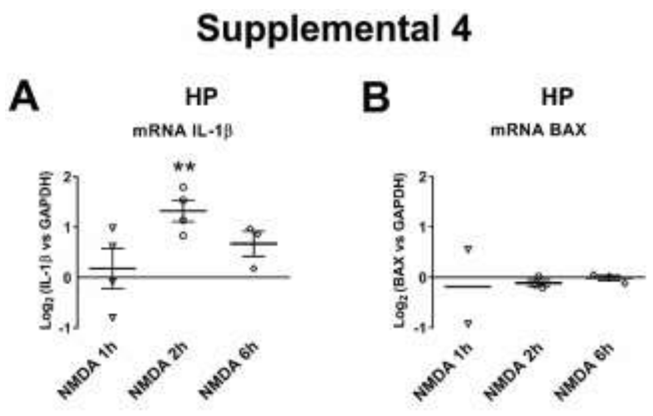


**Supplemental Figure S4. Time course of IL-1β mRNA upregulation in response to NMDA in hippocampal cultures. A) and B)** IL-1β mRNA (left) and BAX mRNA (right) measured by quantitative PCR in hippocampal cultures 1, 2 and 6 hours after 100 µM NMDA stimuli. Bar graphs show the mean ± SEM fold change normalized against GAPDH as reference. Data was obtained from 2 to 4 independent hippocampal cultures. Statistical significance was assessed by One-way ANOVA followed by Bonferroni post-test. **$p<0.01$.

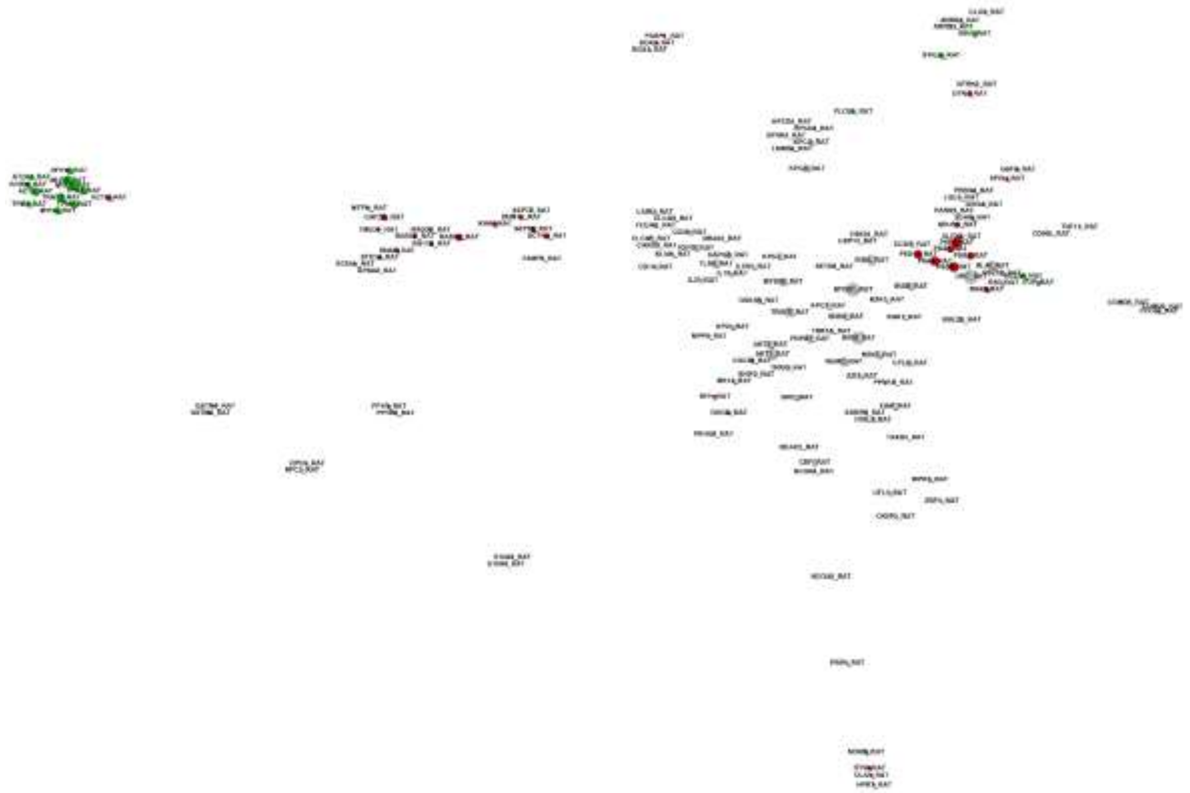

**Supplemental Figure S5**. The annotated Gene Ontology terms of biological processes were retrieved for the identified proteins that were exclusively detected in cortical (red) or hippocampal (green) proteomes and NF-κB related proteins (grey). In Figure 2 G the proteins are depicted as nodes with shared biological processes shown as edges connecting the nodes, forming a graph. To better display the similarity between the different protein groups a force field embedder was used to calculate a layout, thus depicting similar proteins closer to each other.

#### SUPPLEMENTAL TABLES

#### SUPPLEMENTAL TABLE 1

| **Figure 1C** | **Mean ± SE** | **n** | **p value** | | |
|---|---|---|---|---|---|
| HP Control vs HP 30 µM NMDA | 1.00 vs 1.84 ± 0.37 | 5 | > 0.05 | | |
| HP Control vs HP 100 µM NMDA | 1.00 vs 2.07 ± 0.16 | 5 | > 0.001 | | |
| CX Control vs CX 30 µM NMDA | 1.00 vs 1.14 ± 0.17 | 9 | n.s | | |
| CX Control vs CX 100 µM NMDA | 1.00 vs 1.26 ± 0.15 | 8 | n.s | | |
| **Figure 1D** | **Mean ± SE** | **n** | **p value** | | |
| HP Control vs HP 30 µM NMDA | 1.00 vs 0.75 ± 0.16 | 4 | n.s | | |
| HP Control vs HP 100 µM NMDA | 1.00 vs 1.02 ± 0.21 | 4 | n.s | | |
| CX Control vs CX 30 µM NMDA | 1.00 vs 0.79 ± 0.03 | 3 | n.s | | |
| CX Control vs CX 100 µM NMDA | 1.00 vs 0.67 | 2 | n.s | | |
| **Figure 1E** | **Mean ± SE** | **n** | **p value** | **ANOVA Table** | **SS** |
| CX Control vs CX 30 µM NMDA | 1.00 ± 0.05 vs 1.03 ± 0.09 | 12 and 7 | n.s | Between columns<br>Within columns<br>Total | 0,2666<br>1,771<br>2,038 |
| CX Control vs CX 100 µM NMDA | 1.00 ± 0.05 vs 1.20 ± 0.09 | 12 | n.s | | |
| CX 30 µM NMDA vs CX 100 µM NMDA | 1.03 ± 0.09 vs 1.20 ± 0.09 | 7 and 12 | n.s | | |
| **Figure 1F** | **Mean ± SE** | **n** | **p value** | **ANOVA Table** | **SS** |
| HP Control vs HP 30 µM NMDA | 1.00 ± 0.06 vs 1.49 ± 0.13 | 6 | n.s | Between columns<br>Within columns<br>Total | 2,652<br>1,555<br>4,207 |
| HP Control vs HP 100 µM NMDA | 1.00 ± 0.06 vs 1.94 ± 0.18 | 6 | > 0.001 | | |
| H HP 30 µM NMDA vs HP 100 µM NMDA | 1.49 ± 0.13 vs 1.94 ± 0.18 | 6 | n.s | | |
| **Figure 2A** | **Mean ± SE** | **n** | **p value** | **ANOVA Table** | **SS** |
| CX Control vs CX 2 µM Ro 106-9920 | 81.15 ± 1.68 vs 77.87 ± 1.74 | 5 | ns | Between columns<br>Within columns<br>Total | 434,1<br>343,9<br>778,0 |
| CX Control vs CX 5 µM Ro 106-9920 | 81.15 ± 1.68 vs 71.97 ± 2.79 | 5 | ns | | |
| CX Control vs CX 10 µM Ro 106-9920 | 81.15 ± 1.68 vs 68.70 ± 2.17 | 5 | > 0.01 | | |
| HP Control vs HP 2 µM Ro 106-9920 | 82.52 ± 0.90 vs 76.59 ± 1.46 | 5 | ns | Between columns<br>Within columns<br>Total | 451,9<br>171,9<br>623,8 |
| HP Control vs HP 5 µM Ro 106-9920 | 82.52 ± 0.90 vs 75.63 ± 1.39 | 5 | > 0.05 | | |
| HP Control vs HP 10 µM Ro 106-9920 | 82.52 ± 0.90 vs 69.12 ± 1.93 | 5 | > 0.001 | | |
| **Figure 2B** | **Mean ± SE** | **n** | **p value** | **ANOVA Table** | **SS** |

| | | | | | |
|---|---|---|---|---|---|
| CX Control vs CX 30 µM NMDA | 74.26 ± 2.47 vs 75.99 ± 2.53 | 4 | n.s | Between columns | 143,1 |
| CX Control vs CX Ro 106-9920 + 30 µM NMDA | 74.26 ± 2.47 vs 67,96 ± 3.60 | 4 | n.s | Within columns | 305,8 |
| CX 30 µM NMDA vs CX Ro 106-9920 + 30 µM NMDA | 75.99 ± 2.53 vs 67,96 ± 3.60 | 4 | n.s | Total | 448,9 |
| HP Control vs HP 30 µM NMDA | 75.54 ± 1.07 vs 66.53 ± 3.12 | 4 | n.s | Between columns | 162,8 |
| HP Control vs HP Ro 106-9920 + 30 µM NMDA | 75.54 ± 1.07 vs 71,54 ± 3.02 | 4 | n.s | Within columns | 239,8 |
| HP 30 µM NMDA vs HP Ro 106-9920 + 30 µM NMDA | 66.53 ± 3.12 vs 71,54 ± 3.02 | 4 | n.s | Total | 402,6 |
| **Figure 2C** | **Mean ± SE** | **n** | **p value** | **ANOVA Table** | **SS** |
| CX Control vs CX 100 µM NMDA | 75.83 ± 1.52 vs 72.60 ± 3.38 | 4 | n.s | Between columns | 776,1 |
| CX Control vs CX Ro 106-9920 + 100 µM NMDA | 75.83 ± 1.52 vs 57.39 ± 5.30 | 4 | > 0.05 | Within columns | 501,3 |
| CX 100 µM NMDA vs CX Ro 106-9920 + 100 µM NMDA | 72.60 ± 3.38 vs 57.39 ± 5.30 | 4 | n.s | Total | 1277 |
| HP Control vs HP 100 µM NMDA | 79.81 ± 0.39 vs 57.54 ± 3.34 | 4 | > 0.01 | Between columns | 1049 |
| HP Control vs HP Ro 106-9920 + 100 µM NMDA | 79.81 ± 0.39 vs 73.30 ± 4.58 | 4 | n.s | Within columns | 387,6 |
| HP 100 µM NMDA vs HP Ro 106-9920 + 100 µM NMDA | 57.54 ± 3.34 vs 73.30 ± 4.58 | 4 | > 0.05 | Total | 1437 |
| **Figure 3** | **Mean ± SE** | **n** | **p value** | | |
| CX Control vs CX 30 µM NMDA | 1.00 vs 3.16 ± 0.49 | 3 | > 0.01 | | |
| HP Control vs HP 30 µM NMDA | 1.00 vs 0.20 ± 0.02 | 3 | > 0.001 | | |
| CX 30 µM NMDA vs HP 30 µM NMDA | 3.16 ± 0.49 vs 0.20 ± 0.02 | 3 | > 0.001 | | |
| **Figure 4B** | **Mean ± SE** | **n** | **p value** | **ANOVA Table** | **SS** |
| CX control vs CX sheNOS | 0.01 ± 0.0011 vs 0.0046 ± 0.0014 | 7 and 5 | > 0.05 | Between columns | 0,0001098 |
| CX shSC vs CX sheNOS | 0.0107 ± 0.0004 vs 0.0046 ± 0.0014 | 4 and 5 | > 0.05 | Within columns | 0,00008698 |
| CX control vs CX shSC | 0.01 ± 0.0011 vs 0.0107 ± 0.0004 | 7 and 4 | n.s | Total | 0,0001968 |
| **Figure 4D** | **Mean ± SE** | **n** | **p value** | | |
| CX shSC vs CX sheNOS (p65) | 1.00 ± 0.04 vs 0.39 ± 0.14 | 6 | > 0.01 | | |
| HP shSC vs HP sheNOS (p65) | 1.00 ± 0.04 vs 0.42 ± 0.21 | 4 | > 0.05 | | |
| CX shSC vs CX sheNOS (Tubulin 1A) | 1.00 ± 0.03 vs 0.27 ± 0.17 | 4 | > 0.01 | | |
| HP shSC vs HP sheNOS (Tubulin 1A) | 1.00 ± 0.07 vs 0.22 ±0.09 | 4 | > 0.001 | | |
| **Figure 5A** | **Mean ± SE** | **n** | **p value** | **ANOVA Table** | **SS** |
| CX Control vs CX LNIO | 1.00 ± 0.09 vs 1.12 ± 0.12 | 6 | n.s | Between columns | 0,1011 |
| CX Control vs CX 100 µM NMDA | 1.00 ± 0.09 vs 1.11 ± 0.14 | 6 | n.s | Within columns | 2,267 |

| | | | | | |
|---|---|---|---|---|---|
| CX control vs CX LNIO + 100 μM NMDA | 1.00 ± 0.09 vs 1.18 ± 0.18 | 6 | n.s | Total | 2,369 |
| CX LNIO vs CX 100 μM NMDA | 1.12 ± 0.12 vs 1.11 ± 0.14 | 6 | n.s | | |
| CX LNIO vs CX LNIO + 100 μM NMDA | 1.12 ± 0.12 vs 1.18 ± 0.18 | 6 | n.s | | |
| CX 100 μM NMDA vs CX LNIO + 100 μM NMDA | 1.11 ± 0.14 vs 1.18 ± 0.18 | 6 | n.s | | |
| **Figure 5B** | **Mean ± SE** | **n** | **p value** | **ANOVA Table** | **SS** |
| HP Control vs HP LNIO | 1.00 ± 0.05 vs 1.21 ± 0.28 | 5 | n.s | Between columns<br>Within columns<br>Total | 4,148<br>2,694<br>6,842 |
| HP Control vs HP 100 μM NMDA | 1.00 ± 0.05 vs 2.06 ± 0.17 | 5 | > 0.01 | | |
| HP control vs HP LNIO + 100 μM NMDA | 1.00 ± 0.04 vs 1.93 ± 0.17 | 5 | > 0.05 | | |
| HP LNIO vs HP 100 μM NMDA | 1.21 ± 0.28 vs 2.06 ± 0.17 | 5 | > 0.05 | | |
| HP LNIO vs HP LNIO + 100 μM NMDA | 1.21 ± 0.28 vs 1.93 ± 0.17 | 5 | n.s | | |
| HP 100 μM NMDA vs HP LNIO + 100 μM NMDA | 1.93 ± 0.17 vs 2.06 ± 0.17 | 5 | n.s | | |
| **Figure 5C** | **Mean ± SE** | **n** | **p value** | **ANOVA Table** | **SS** |
| CX Control vs CX LNIO | 1.00 ± 0.05 vs 0.98 ± 0.13 | 10 and 5 | n.s | Between columns<br>Within columns<br>Total | 1,572<br>2,104<br>3,675 |
| CX Control vs CX LNIO + 30 μM NMDA | 1.00 ± 0.05 vs 1.24 ± 0.11 | 10 and 5 | n.s | | |
| CX Control vs CX LNIO + 100 μM NMDA | 1.00 ± 0.05 vs 1.52 ± 0.13 | 10 and 9 | > 0.01 | | |
| CX LNIO vs CX LNIO + 30 μM NMDA | 0.98 ± 0.13 vs 1.24 ± 0.11 | 5 | n.s | | |
| CX LNIO vs CX LNIO + 100 μM NMDA | 0.98 ± 0.13 vs 1.52 ± 0.13 | 5 and 9 | > 0.01 | | |
| CX LNIO + 30 μM NMDA vs CX LNIO + 100 μM NMDA | 1.24 ± 0.11 vs 1.52 ± 0.13 | 5 and 9 | n.s | | |
| **Figure 5D** | **Mean ± SE** | **n** | **p value** | **ANOVA Table** | **SS** |
| HP Control vs HP LNIO | 1.00 ± 0.04 vs 1.04 ± 0.16 | 6 and 4 | n.s | Between columns<br>Within columns<br>Total | 1,067<br>0,4511<br>1,518 |
| HP Control vs HP LNIO + 30 μM NMDA | 1.00 ± 0.04 vs 1.27 ± 0.06 | 6 and 4 | n.s | | |
| HP Control vs HP 100 μM NMDA | 1.00 ± 0.04 vs 1.62 ± 0.07 | 6 and 4 | > 0.01 | | |
| HP LNIO vs HP LNIO 30 μM NMDA | 1.04 ± 0.16 vs 1.27 ± 0.06 | 4 | n.s | | |
| HP LNIO vs HP LNIO 100 μM NMDA | 1.04 ± 0.16 vs 1.62 ± 0.07 | 4 | > 0.01 | | |
| HP LNIO + 30 μM NMDA vs HP LNIO + 100 μM NMDA | 1.27 ± 0.06 vs 1.62 ± 0.07 | 4 | n.s | | |
| **Figure 5E** | **Mean ± SE** | **n** | **p value** | **ANOVA Table** | **SS** |
| HP Control vs HP SNAP | 1.00 ± 0.06 vs 1.07 ± 0.11 | 6 and 5 | n.s | Between columns<br>Within columns<br>Total | 2,107<br>1,857<br>3,964 |
| HP Control vs HP 100 μM NMDA | 1.00 ± 0.06 vs 1.76 ± 0.17 | 6 | > 0.01 | | |
| HP Control vs HP SNAP + 100 μM NMDA | 1.00 ± 0.06 vs 1.20 ± 0.14 | 6 | n.s | | |

| | | | | | |
|---|---|---|---|---|---|
| HP SNAP vs HP 100 µM NMDA | 1.07 ± 0.11 vs 1.76 ± 0.17 | 5 and 6 | > 0.01 | | |
| HP SNAP vs HP SNAP + 100 µM NMDA | 1.07 ± 0.11 vs 1.20 ± 0.14 | 5 and 6 | n.s | | |
| HP 100 µM NMDA vs HP SNAP + 100 µM NMDA | 1.76 ± 0.17 vs 1.20 ± 0.14 | 6 | > 0.05 | | |
| **Figure 5F** | **Mean ± SE** | **n** | **p value** | **ANOVA Table** | **SS** |
| HP Control vs HP 100 µM NMDA | 1.00 vs 2.32 ± 0.31 | 8 | > 0.001 | Between columns<br>Within columns<br>Total | SS<br>11,95<br>5,726<br>17,68 |
| HP Control vs HP Ro 106-9920 | 1.00 vs 0.55 ± 0.12 | 8 and 4 | n.s | | |
| HP Control vs HP Ro106-9920 + 100 µM NMDA | 1.00 vs 0.88 ± 0.16 | 8 and 4 | n.s | | |
| HP 100 µM NMDA vs HP Ro 106-9920 | 2.32 ± 0.31 vs 0.55 ± 0.12 | 8 and 4 | > 0.001 | | |
| HP 100 µM NMDA vs HP Ro106-9920 + 100 µM NMDA | 2.32 ± 0.31 vs 0.88 ± 0.16 | 8 and 4 | > 0.001 | | |
| HP Ro106-9920 vs HP Ro106-9920 + 100 µM NMDA | 0.55 ± 0.12 vs 0.88 ± 0.16 | 4 | n.s | | |
| **Figure 5G** | **Mean ± SE** | **n** | **p value** | **ANOVA Table** | **SS** |
| HP Control vs HP 100 µM NMDA | 1.00 vs 2.32 ± 0.31 | 8 | > 0.001 | Between columns<br>Within columns<br>Total | 12,62<br>8,880<br>21,50 |
| HP Control vs HP SNAP | 1.00 vs 0.76 ± 0.13 | 8 and 4 | n.s | | |
| HP Control vs HP SNAP + 100 µM NMDA | 1.00 vs 1.10 ± 0.40 | 8 and 5 | n.s | | |
| HP 100 µM NMDA vs HP SNAP | 2.48 ± 0.31 vs 0.76 ± 0.13 | 8 and 4 | > 0.01 | | |
| HP 100 µM NMDA vs HP SNAP + 100 µM NMDA | 2.48 ± 0.31 vs 1.10 ± 0.40 | 8 and 5 | > 0.01 | | |
| HP SNAP vs HP SNAP + 100 µM NMDA | 0.76 ± 0.13 vs 1.10 ± 0.40 | 4 and 5 | n.s | | |
| **Figure 6D** | **Mean ± SE** | **n** | **p value** | | |
| CX Control vs CX 30 µM NMDA (GluN2A) | 1.00 vs 1.86 ± 0.33 | 5 | > 0.05 | | |
| HP Control vs HP 30 µM NMDA (GluN2A) | 1.00 vs 1.76 ± 0.15 | 5 | > 0.01 | | |
| CX 30 µM NMDA vs HP 30 µM NMDA (GluN2A) | 1.86 ± 0.33 vs 1.76 ± 0.15 | 5 | n.s | | |
| CX Control vs CX 30 µM NMDA (PSD95) | 1.00 vs 2.02 ± 0.01 | 3 | > 0.001 | | |
| HP Control vs HP 30 µM NMDA (PSD95) | 1.00 vs 1.70 ± 0.13 | 3 | > 0.01 | | |
| CX 30 µM NMDA vs HP 30 µM NMDA (PSD95) | 2.02 ± 0.01 vs 1.70 ± 0.13 | 3 | n.s | | |
| CX Control vs CX 30 µM NMDA (SAPAP4) | 1.00 vs 1.58 ± 0.09 | 4 | > 0.001 | | |
| HP Control vs HP 30 µM NMDA (SAPAP4) | 1.00 vs 1.15 ± 0.06 | 3 | n.s | | |
| CX 30 µM NMDA vs HP 30 µM NMDA (SAPAP4) | 1.58 ± 0.09 vs 1.15 ± 0.06 | 3 and 4 | > 0.05 | | |
| **Figure S1D** | **Mean ± SE** | **n** | **p value** | **ANOVA Table** | **SS** |
| CX Control vs CX 30 µM NMDA | 72.58 ± 2.61 vs 68.94 ± 1.23 | 3 | n.s | Between columns | 185,1 |

| | | | | | |
|---|---|---|---|---|---|
| CX Control vs CX 60 µM NMDA | 72.58 ± 2.61 vs 63.26 ± 3.75 | 3 | n.s | Within columns | 911,3 |
| CX Control vs CX 100 µM NMDA | 72.58 ± 2.61 vs 65.04 ± 4.66 | 3 | n.s | Total | 1096 |
| CX 30 µM NMDA vs CX 60 µM NMDA | 68.94 ± 1.23 vs 63.26 ± 3.75 | 3 | n.s | | |
| CX 30 µM NMDA vs CX 100 µM NMDA | 68.94 ± 1.23 vs 65.04 ± 4.66 | 3 | n.s | | |
| CX 60 µM NMDA vs CX 100 µM NMDA | 63.26 ± 3.75 vs 65.04 ± 4.66 | 3 | n.s | | |
| HP Control vs HP 30 µM NMDA | 73.90 ± 3.96 vs 70.61 ± 4.14 | 4 | n.s | Between columns | 1049 |
| HP Control vs HP 60 µM NMDA | 73.90 ± 3.96 vs 53.25 ± 5.15 | 4 | > 0.05 | Within columns<br>Total | 387,6<br>1437 |
| HP Control vs HP 100 µM NMDA | 73.90 ± 3.96 vs 40.12 ± 4.53 | 4 | > 0.01 | | |
| HP 30 µM NMDA vs HP 60 µM NMDA | 70.61 ± 4.14 vs 53.25 ± 5.15 | 4 | n.s | | |
| HP 30 µM NMDA vs HP 100 µM NMDA | 70.61 ± 4.14 vs 40.12 ± 4.53 | 4 | > 0.01 | | |
| HP 60 µM NMDA vs HP 100 µM NMDA | 53.25 ± 5.15 vs 40.12 ± 4.53 | 4 | n.s | | |
| **Figure S1E** | **Mean ± SE** | **n** | **p value** | | |
| Control vs NMDA | 1.00 ± 0.13 vs 18.62 ± 2.79 | 13 | > 0.001 | | |
| **Figure S2C** | **Mean ± SE** | **n** | **p value** | | |
| HP Control vs HP NMDA (Neurons) | 1.00 vs 1.75 ± 0.26 | 3 | > 0.05 | | |
| HP Control vs HP NMDA (Astrocytes) | 1.00 vs 1.48 ± 0.17 | 3 | > 0.05 | | |
| **Figure S3A** | **Mean ± SE** | **n** | **p value** | **ANOVA Table** | **SS** |
| CX Control vs CX LNIO | 1.00 ± 0.09 vs 1.10 ± 0.13 | 5 | n.s | Between columns | 0,3149 |
| CX Control vs CX 100 µM NMDA | 1.00 ± 0.09 vs 1.08 ± 0.15 | 5 | n.s | Within columns<br>Total | 1,309<br>1,624 |
| CX Control vs CX LNIO + 100 µM NMDA | 1.00 ± 0.09 vs 0.78 ± 0.14 | 5 | n.s | | |
| CX LNIO vs CX 100 µM NMDA | 1.10 ± 0.13 vs 1.08 ± 0.15 | 5 | n.s | | |
| CX LNIO vs CX LNIO + 100 µM NMDA | 1.10 ± 0.13 vs 0.78 ± 0.14 | 5 | n.s | | |
| CX 100 µM NMDA vs CX LNIO + 100 µM NMDA | 1.08 ± 0.15 vs 0.78 ± 0.14 | 5 | n.s | | |
| **Figure S3B** | **Mean ± SE** | **n** | **p value** | **ANOVA Table** | **SS** |
| HP Control vs HP LNIO | 1.00 ± 0.08 vs 0.94 ± 0.15 | 4 | n.s | Between columns | 0,009124 |
| HP Control vs HP 100 µM NMDA | 1.00 ± 0.08 vs 0.96 ± 0.14 | 4 | n.s | Within columns<br>Total | 0,7017<br>0,7108 |
| HP Control vs HP LNIO + 100 µM NMDA | 1.00 ± 0.08 vs 0.94 ± 0.09 | 4 | n.s | | |
| HP LNIO vs HP 100 µM NMDA | 0.94 ± 0.15 vs 0.96 ± 0.14 | 4 | n.s | | |
| HP LNIO vs HP LNIO + 100 µM NMDA | 0.94 ± 0.15 vs 0.94 ± 0.09 | 4 | n.s | | |

| | | | | | |
|---|---|---|---|---|---|
| HP 100 µM NMDA vs HP LNIO + 100 µM NMDA | 0.96 ± 0.14 vs 0.94 ± 0.09 | 4 | n.s | | |
| **Figure S3C** | **Mean ± SE** | **n** | **p value** | **ANOVA Table** | **SS** |
| CX Control vs CX LNIO | 1.00 ± 0.19 vs 0.71 ± 0.12 | 6 | n.s | Between columns<br>Within columns<br>Total | 0,5933<br>2,799<br>3,392 |
| CX Control vs CX 100 µM NMDA | 1.00 ± 0.19 vs 0.88 ± 0.19 | 6 | n.s | | |
| CX Control vs CX LNIO + 100 µM NMDA | 1.00 ± 0.19 vs 0.59 ± 0.08 | 6 | n.s | | |
| CX LNIO vs CX 100 µM NMDA | 0.71 ± 0.12 vs 0.88 ± 0.19 | 6 | n.s | | |
| CX LNIO vs CX LNIO + 100 µM NMDA | 0.71 ± 0.12 vs 0.59 ± 0.08 | 6 | n.s | | |
| CX 100 µM NMDA vs CX LNIO + 100 µM NMDA | 0.88 ± 0.19 vs 0.59 ± 0.08 | 6 | n.s | | |
| **Figure S3D** | **Mean ± SE** | **n** | **p value** | **ANOVA Table** | **SS** |
| HP Control vs HP LNIO | 1.00 ± 0.19 vs 0.98 ± 0.20 | 5 | n.s | Between columns<br>Within columns<br>Total | 1,445<br>7,858<br>9,303 |
| HP Control vs HP 100 µM NMDA | 1.00 ± 0.19 vs 1.64 ± 0.51 | 5 | n.s | | |
| HP Control vs HP LNIO + 100 µM NMDA | 1.00 ± 0.19 vs 1.10 ± 0.23 | 5 | n.s | | |
| HP LNIO vs HP 100 µM NMDA | 0.98 ± 0.19 vs 1.64 ± 0.51 | 5 | n.s | | |
| HP LNIO vs HP LNIO + 100 µM NMDA | 0.98 ± 0.19 vs 1.10 ± 0.23 | 5 | n.s | | |
| HP 100 µM NMDA vs HP LNIO + 100 µM NMDA | 1.64 ± 0.51 vs 1.10 ± 0.23 | 5 | n.s | | |
| **Figure S4A** | **Mean ± SE** | **n** | **p value** | **ANOVA Table** | **SS** |
| HP Control vs 100 µM NMDA 1h | 1.00 vs 1.25 ± 0.31 | 4 | n.s | Between columns<br>Within columns<br>Total | 5,784<br>3,253<br>9,037 |
| HP Control vs 100 µM NMDA 2h | 1.00 vs 2.58 ± 0.38 | 4 | > 0.01 | | |
| HP Control vs 100 µM NMDA 6h | 1.00 vs 1.64 ± 0.26 | 4 | n.s | | |
| **Figure S4B** | **Mean ± SE** | **n** | **p value** | **ANOVA Table** | **SS** |
| HP Control vs 100 µM NMDA 1h | 1.00 vs 1.00 ± 0.47 | 2 | n.s | Between columns<br>Within columns<br>Total | 0,01717<br>0,4623<br>0,4795 |
| HP Control vs 100 µM NMDA 2h | 1.00 vs 0.91 ± 0.02 | 3 | n.s | | |
| HP Control vs 100 µM NMDA 6h | 1.00 vs 0.99 ± 0.04 | 3 | n.s | | |

**Supplemental Table 1. Statistics of all experiments**

#### SUPPLEMENTAL TABLE 2

| | | Treatment | | | |
|---|---|---|---|---|---|
| | | Hippocampal Cultures | | Cortical Cultures | |
| **Protein name** | **Entry name** | **HP Control** | **HP NMDA** | **CX Control** | **CX NMDA** |
| **n** | | 6 | 5 | 6 | 6 |
| 14-3-3 protein epsilon | 1433E_RAT | 5 | 3 | 2 | 3 |
| 14-3-3 protein eta | 1433F_RAT | 5 | 3 | 2 | 2 |
| 14-3-3 protein gamma | 1433G_RAT | 6 | 4 | 2 | 2 |
| 14-3-3 protein theta | 1433T_RAT | 6 | 4 | 2 | 2 |
| 14-3-3 protein zeta/delta | 1433Z_RAT | 0 | 0 | 6 | 6 |
| 1-phosphatidylinositol-4,5-bisphosphate phosphodiesterase beta-1 | PLCB1_RAT | 0 | 1 | 0 | 0 |
| 26S protease regulatory subunit 4 | PRS4_RAT | 1 | 1 | 0 | 0 |
| 26S protease regulatory subunit 6A | PRS6A_RAT | 0 | 0 | 2 | 2 |
| 26S protease regulatory subunit 6B | PRS6B_RAT | 0 | 0 | 3 | 3 |
| 26S protease regulatory subunit 7 | PRS7_RAT | 1 | 2 | 0 | 0 |
| 26S protease regulatory subunit 8 | PRS8_RAT | 0 | 0 | 1 | 2 |
| 26S proteasome non-ATPase regulatory subunit 11 | PSD11_RAT | 0 | 0 | 1 | 1 |
| 26S proteasome non-ATPase regulatory subunit 13 | PSD13_RAT | 0 | 0 | 0 | 1 |
| 26S proteasome non-ATPase regulatory subunit 2 | PSMD2_RAT | 1 | 3 | 2 | 3 |
| 2-oxoglutarate dehydrogenase, mitochondrial | ODO1_RAT | 0 | 0 | 1 | 2 |

| | | | | | |
|---|---|---|---|---|---|
| 40S ribosomal protein S17 | RS17_RAT | 1 | 0 | 0 | 0 |
| 40S ribosomal protein S3 | RS3_RAT | 0 | 0 | 2 | 3 |
| 40S ribosomal protein S4, X isoform | RS4X_RAT | 0 | 0 | 0 | 2 |
| 40S ribosomal protein S8 | RS8_RAT | 0 | 0 | 1 | 0 |
| 40S ribosomal protein SA | RSSA_RAT | 3 | 2 | 3 | 3 |
| 4-aminobutyrate aminotransferase, mitochondrial | GABT_RAT | 0 | 0 | 2 | 3 |
| 4F2 cell-surface antigen heavy chain | 4F2_RAT | 4 | 4 | 2 | 3 |
| 4-trimethylaminobutyraldehyde dehydrogenase | AL9A1_RAT | 0 | 0 | 1 | 2 |
| 60 kDa heat shock protein, mitochondrial | CH60_RAT | 4 | 4 | 3 | 3 |
| 60S acidic ribosomal protein P0 | RLA0_RAT | 0 | 0 | 3 | 3 |
| 60S ribosomal protein L19 | RL19_RAT | 0 | 0 | 1 | 0 |
| 60S ribosomal protein L23a | RL23A_RAT | 0 | 1 | 0 | 0 |
| 60S ribosomal protein L3 | RL3_RAT | 0 | 0 | 3 | 2 |
| 60S ribosomal protein L4 | RL4_RAT | 0 | 0 | 2 | 2 |
| 60S ribosomal protein L5 | RL5_RAT | 0 | 0 | 1 | 0 |
| 6-phosphofructokinase | K6PP_RAT | 1 | 3 | 0 | 1 |
| 6-phosphofructokinase, liver type | PFKAL_RAT | 4 | 3 | 1 | 2 |
| 6-phosphofructokinase, muscle type | PFKAM_RAT | 0 | 0 | 2 | 2 |
| 6-phosphogluconate dehydrogenase, decarboxylating | 6PGD_RAT | 4 | 2 | 3 | 2 |
| 6-phosphogluconolactonase | 6PGL_RAT | 0 | 0 | 0 | 1 |
| 78 kDa glucose-regulated protein | GRP78_RAT | 0 | 0 | 1 | 3 |

| | | | | | |
|---|---|---|---|---|---|
| 7-dehydrocholesterol reductase | DHCR7_RAT | 2 | 0 | 1 | 0 |
| Acetyl-CoA acetyltransferase, cytosolic | THIC_RAT | 0 | 0 | 2 | 3 |
| Acetyl-CoA acetyltransferase, mitochondrial | THIL_RAT | 0 | 0 | 1 | 3 |
| Aconitate hydratase, mitochondrial | ACON_RAT | 5 | 2 | 3 | 3 |
| Actin cytoplasmic 2 | ACTG_RAT | 0 | 0 | 1 | 2 |
| Actin, alpha cardiac muscle 1 | ACTC_RAT | 0 | 0 | 0 | 2 |
| Actin, alpha skeletal muscle | ACTS_RAT | 0 | 1 | 0 | 0 |
| Actin-related protein 2 | ARP2_RAT | 0 | 0 | 2 | 2 |
| Actin-related protein 2/3 complex subunit 1A | ARC1A_RAT | 0 | 0 | 1 | 3 |
| Adenosylhomocysteinase | SAHH_RAT | 4 | 2 | 1 | 0 |
| Adenylate kinase isoenzyme 1 | KAD1_RAT | 0 | 1 | 0 | 0 |
| Adenylyl cyclase-associated protein 1 | CAP1_RAT | 2 | 3 | 3 | 3 |
| ADP/ATP translocase 1 | ADT1_RAT | 0 | 0 | 3 | 3 |
| ADP/ATP translocase 2 | ADT2_RAT | 2 | 2 | 1 | 1 |
| Alanine--tRNA ligase, cytoplasmic | SYAC_RAT | 0 | 0 | 3 | 3 |
| Alcohol dehydrogenase [NADP(+)] | ADHX_RAT | 0 | 0 | 1 | 2 |
| Aldehyde dehydrogenase, mitochondrial | ALDH2_RAT | 2 | 2 | 1 | 3 |
| Alpha-actinin-4 | ACTN4_RAT | 0 | 0 | 1 | 2 |
| Alpha-aminoadipic semialdehyde dehydrogenase | AL7A1_RAT | 1 | 1 | 1 | 0 |
| Alpha-centractin | ACTZ_RAT | 0 | 0 | 3 | 3 |
| Alpha-tubulin N-acetyltransferase 1 | ATAT_RAT | 0 | 0 | 2 | 0 |

| | | | | | |
|---|---|---|---|---|---|
| Amine oxidase [flavin-containing] B | AOFB_RAT | 4 | 2 | 0 | 0 |
| Amphiphysin | AMPH_RAT | 0 | 1 | 0 | 0 |
| Anionic trypsin-1 | TRY1_RAT | 0 | 0 | 0 | 1 |
| AP-2 complex subunit alpha-2 | AP2A2_RAT | 0 | 0 | 3 | 3 |
| AP-2 complex subunit beta | AP2B1_RAT | 0 | 0 | 3 | 3 |
| AP2-associated protein kinase 1 | AAK1_RAT | 0 | 0 | 1 | 1 |
| ARF GTPase-activating protein GIT1 | APOE_RAT | 0 | 0 | 1 | 1 |
| ArfGAP with GTPase domain, ankyrin repeat and PH domain 3 | ARGI1_RAT | 0 | 0 | 1 | 0 |
| Arginine--tRNA ligase, cytoplasmic | SYRC_RAT | 0 | 0 | 1 | 0 |
| Armadillo repeat-containing protein 10 | ARM10_RAT | 0 | 0 | 0 | 1 |
| Asparaginyl-tRNA synthetase | AATC_RAT | 0 | 0 | 0 | 3 |
| Aspartate aminotransferase, mitochondrial | AATM_RAT | 4 | 4 | 3 | 3 |
| Aspartate--tRNA ligase, cytoplasmic | SYDC_RAT | 0 | 0 | 1 | 0 |
| ATP synthase F(0) complex subunit B1, mitochondrial | AT5F1_RAT | 0 | 0 | 1 | 1 |
| ATP synthase subunit gamma, mitochondrial | ATPG_RAT | 3 | 4 | 1 | 2 |
| Beta-soluble NSF attachment protein | SNAB_RAT | 0 | 0 | 0 | 1 |
| Bleomycin hydrolase | BLMH_RAT | 0 | 0 | 1 | 1 |
| Brain acid soluble protein 1 | BASP1_RAT | 0 | 2 | 0 | 0 |
| Calcium/calmodulin-dependent protein kinase type II subunit alpha | KCC2A_RAT | 0 | 0 | 3 | 3 |
| Calnexin | CALX_RAT | 0 | 0 | 2 | 3 |

| | | | | | |
|---|---|---|---|---|---|
| Calpain-2 catalytic subunit | CAN2_RAT | 0 | 0 | 0 | 2 |
| Calreticulin | CALR_RAT | 0 | 0 | 2 | 3 |
| CaM kinase-like vesicle-associated protein | CAMKV_RAT | 0 | 0 | 0 | 1 |
| cAMP-dependent protein kinase catalytic subunit beta | KAPCB_RAT | 0 | 0 | 2 | 3 |
| Casein kinase II subunit alpha | CSK21_RAT | 0 | 0 | 0 | 2 |
| Catalase | CATA_RAT | 0 | 0 | 1 | 1 |
| Catenin beta-1 | CTNB1_RAT | 0 | 0 | 2 | 3 |
| Cathepsin D | CATD_RAT | 4 | 1 | 2 | 2 |
| Cell adhesion molecule 3 | CADM3_RAT | 0 | 0 | 1 | 1 |
| cGMP-dependent 3',5'-cyclic phosphodiesterase | PDE2A_RAT | 0 | 0 | 1 | 2 |
| Citrate synthase, mitochondrial | CISY_RAT | 4 | 3 | 0 | 0 |
| Clathrin coat assembly protein AP180 | AP180_RAT | 1 | 3 | 0 | 0 |
| CLIP-associating protein 2 | CLAP2_RAT | 0 | 0 | 2 | 3 |
| Coatomer subunit beta | COPB_RAT | 0 | 0 | 1 | 0 |
| Coatomer subunit gamma-1 | COPG1_RAT | 0 | 0 | 2 | 3 |
| Contactin-1 | CNTN1_RAT | 0 | 0 | 3 | 3 |
| Coronin-1A | COR1A_RAT | 3 | 2 | 3 | 3 |
| Creatine kinase M-type | KCRM_RAT | 0 | 1 | 0 | 0 |
| Cullin-3 | CUL3_RAT | 0 | 0 | 1 | 0 |
| Cullin-associated NEDD8-dissociated protein 1 | CAND1_RAT | 0 | 0 | 2 | 3 |
| Cytochrome b-c1 complex subunit 1, mitochondrial | QCR1_RAT | 0 | 0 | 3 | 3 |

| | | | | | |
|---|---|---|---|---|---|
| Cytochrome b-c1 complex subunit 2, mitochondrial | QCR2_RAT | 2 | 4 | 2 | 3 |
| Cytosolic non-specific dipeptidase | CNDP2_RAT | 0 | 0 | 2 | 0 |
| D-3-phosphoglycerate dehydrogenase | SERA_RAT | 3 | 4 | 3 | 3 |
| D-beta-hydroxybutyrate dehydrogenase, mitochondrial | BDH_RAT | 0 | 1 | 0 | 0 |
| Dihydrolipoyl dehydrogenase, mitochondrial | DLDH_RAT | 0 | 0 | 1 | 1 |
| Dihydropyrimidinase-related protein 1 | DPYL1_RAT | 0 | 0 | 3 | 3 |
| Dihydropyrimidinase-related protein 4 (Fragment) | DPYL4_RAT | 3 | 3 | 2 | 1 |
| Dihydropyrimidinase-related protein 5 | DPYL5_RAT | 5 | 4 | 1 | 1 |
| Dipeptidyl peptidase 2 | DPP2_RAT | 3 | 2 | 1 | 0 |
| Dipeptidyl peptidase 3 | DPP3_RAT | 0 | 0 | 1 | 3 |
| Discs, large (Drosophila) homolog-associated protein 4 | DLGP4_RAT | 0 | 0 | 0 | 4 |
| DnaJ homolog subfamily A member 1 | DNJA1_RAT | 2 | 3 | 0 | 0 |
| DnaJ homolog subfamily A member 2 | DNJA2_RAT | 0 | 0 | 1 | 3 |
| Dolichyl-diphosphooligosaccharide--protein glycosyltransferase subunit 1 | RPN1_RAT | 1 | 3 | 0 | 0 |
| Dynactin subunit 1 | DCTN1_RAT | 0 | 0 | 0 | 2 |
| Dynamin-1 | DYN1_RAT | 4 | 3 | 0 | 0 |
| Dynamin-like 120 kDa protein, mitochondrial | OPA1_RAT | 0 | 0 | 2 | 3 |
| EH domain-containing protein 1 | EHD1_RAT | 0 | 0 | 1 | 2 |
| EH domain-containing protein 3 | EHD3_RAT | 1 | 0 | 1 | 1 |
| Electron transfer flavoprotein subunit alpha, mitochondrial | ETFA_RAT | 3 | 2 | 0 | 0 |
| Elongation factor 1-alpha 2 | EF1A2_RAT | 0 | 1 | 0 | 1 |

| | | | | | |
|---|---|---|---|---|---|
| Elongation factor 1-delta | EF1D_RAT | 3 | 2 | 2 | 3 |
| Elongation factor 1-gamma | EF1G_RAT | 0 | 0 | 3 | 3 |
| Elongation factor 2 | EF2_RAT | 0 | 0 | 3 | 3 |
| Elongation factor Tu, mitochondrial | EFTU_RAT | 0 | 0 | 2 | 3 |
| Endoplasmin | ENPL_RAT | 0 | 0 | 2 | 2 |
| Epsin-1 | EPN1_RAT | 0 | 0 | 1 | 0 |
| ES1 protein homolog, mitochondrial | ES1_RAT | 0 | 0 | 0 | 1 |
| Eukaryotic initiation factor 4A-II | IF4A2_RAT | 0 | 0 | 3 | 3 |
| Eukaryotic translation initiation factor 2 subunit 3 | IF2G_RAT | 0 | 0 | 2 | 1 |
| Eukaryotic translation initiation factor 3 subunit A | EIF3A_RAT | 0 | 0 | 1 | 2 |
| Eukaryotic translation initiation factor 3 subunit B | EIF3B_RAT | 0 | 1 | 0 | 1 |
| Eukaryotic translation initiation factor 5B | IF2P_RAT | 0 | 1 | 0 | 0 |
| Excitatory amino acid transporter 1 | EAA1_RAT | 3 | 3 | 0 | 0 |
| Exocyst complex component 4 | EXOC4_RAT | 0 | 0 | 1 | 0 |
| Exportin-1 | EXPO1_RAT | 0 | 0 | 1 | 3 |
| Ezrin | EZRI_RAT | 0 | 0 | 1 | 0 |
| F-actin-capping protein subunit alpha-2 | CAZA2_RAT | 0 | 0 | 2 | 2 |
| F-actin-capping protein subunit beta | CAPZB_RAT | 0 | 0 | 0 | 1 |
| Fascin | FSCN1_RAT | 5 | 3 | 3 | 3 |
| Fatty acid synthase | FAS_RAT | 1 | 3 | 3 | 3 |
| Fatty acid-binding protein, brain | FABP7_RAT | 1 | 0 | 0 | 0 |

| | | | | | |
|---|---|---|---|---|---|
| FERM, RhoGEF and pleckstrin domain-containing protein 1 | FARP1_RAT | 0 | 0 | 0 | 1 |
| Fructose-bisphosphate aldolase A | ALDOA_RAT | 0 | 0 | 3 | 3 |
| Fructose-bisphosphate aldolase C | ALDOC_RAT | 0 | 0 | 3 | 3 |
| Fumarate hydratase, mitochondrial | FUMH_RAT | 2 | 1 | 0 | 0 |
| Gamma-enolase | ENOG_RAT | 4 | 4 | 3 | 3 |
| General vesicular transport factor p115 | USO1_RAT | 0 | 0 | 1 | 1 |
| Glucose-6-phosphate 1-dehydrogenase | G6PD_RAT | 0 | 0 | 0 | 1 |
| Glucose-6-phosphate isomerase | G6PI_RAT | 4 | 4 | 2 | 3 |
| Glutamate decarboxylase 2 | DCE2_RAT | 0 | 0 | 1 | 0 |
| Glutamate dehydrogenase 1, mitochondrial | DHE3_RAT | 4 | 4 | 3 | 3 |
| Glutaminase kidney isoform, mitochondrial | GLSK_RAT | 1 | 2 | 0 | 1 |
| Glutamine synthetase | GLNA_RAT | 3 | 4 | 0 | 2 |
| Glutathione S-transferase alpha-3 | GSTA3_RAT | 1 | 2 | 0 | 0 |
| Glutathione S-transferase Mu 1 | GSTM1_RAT | 2 | 2 | 0 | 0 |
| Glutathione S-transferase Mu 5 | GSTM5_RAT | 0 | 2 | 0 | 0 |
| Glutathione S-transferase P | GSTP1_RAT | 0 | 0 | 1 | 1 |
| Glutathione S-transferase Yb-3 | GSTM4_RAT | 0 | 0 | 0 | 1 |
| Glyceraldehyde-3-phosphate dehydrogenase | G3P_RAT | 0 | 0 | 2 | 3 |
| Glycine amidinotransferase, mitochondrial | GATM_RAT | 0 | 0 | 2 | 3 |
| Glycine--tRNA ligase | GARS_RAT | 0 | 0 | 1 | 2 |
| Glycogen phosphorylase, brain form (Fragment) | PYGB_RAT | 0 | 2 | 1 | 3 |

| | | | | | |
|---|---|---|---|---|---|
| Glycogen synthase kinase-3 beta | GSK3B_RAT | 0 | 0 | 2 | 3 |
| GPI inositol-deacylase | PGAP1_RAT | 0 | 0 | 1 | 0 |
| GTP-binding nuclear protein Ran | RAN_RAT | 0 | 0 | 2 | 1 |
| Guanine deaminase | GUAD_RAT | 1 | 1 | 0 | 0 |
| Guanine nucleotide-binding protein G(i) subunit alpha-1 | GNAI1_RAT | 0 | 0 | 3 | 3 |
| Guanine nucleotide-binding protein G(i) subunit alpha-2 | GNAI2_RAT | 0 | 0 | 2 | 2 |
| Guanine nucleotide-binding protein G(I)/G(S)/G(T) subunit beta-1 | GBB1_RAT | 4 | 4 | 3 | 3 |
| Guanine nucleotide-binding protein G(I)/G(S)/G(T) subunit beta-2 | GBB2_RAT | 3 | 3 | 2 | 3 |
| Guanine nucleotide-binding protein G(o) subunit alpha | GNAO_RAT | 0 | 0 | 2 | 3 |
| Guanine nucleotide-binding protein G(s) subunit alpha isoforms short | GNAS2_RAT | 0 | 0 | 2 | 3 |
| Guanine nucleotide-binding protein G(z) subunit alpha | GNAZ_RAT | 0 | 0 | 0 | 1 |
| Guanine nucleotide-binding protein subunit alpha-12 | GNA12_RAT | 0 | 0 | 1 | 0 |
| Guanine nucleotide-binding protein subunit beta-2-like 1 | GBLP_RAT | 3 | 3 | 1 | 2 |
| Guanine nucleotide-binding protein-like 1 | GNL1_RAT | 0 | 0 | 1 | 0 |
| Guanylate cyclase soluble subunit alpha-3 | GCYA3_RAT | 0 | 0 | 1 | 0 |
| Heat shock 70 kDa protein 4 | HSP74_RAT | 4 | 2 | 0 | 0 |
| Heat shock cognate 71 kDa | HSP7C_RAT | 0 | 0 | 3 | 3 |
| Heat shock protein 105 kDa | HS105_RAT | 0 | 1 | 0 | 0 |
| Heat shock protein 75 kDa, mitochondrial | TRAP1_RAT | 0 | 0 | 1 | 2 |

| | | | | | |
|---|---|---|---|---|---|
| Heat shock protein HSP 90-alpha | HS90A_RAT | 0 | 0 | 3 | 3 |
| Heat shock protein HSP 90-beta | HS90B_RAT | 0 | 0 | 3 | 3 |
| Heterogeneous nuclear ribonucleoprotein A1 | ROA1_RAT | 0 | 2 | 0 | 0 |
| Heterogeneous nuclear ribonucleoprotein A3 | ROA3_RAT | 0 | 0 | 0 | 1 |
| Heterogeneous nuclear ribonucleoprotein K | HNRPK_RAT | 0 | 0 | 2 | 3 |
| Hexokinase-1 | HXK1_RAT | 5 | 4 | 3 | 3 |
| High mobility group protein B1 | HMGB1_RAT | 0 | 2 | 0 | 0 |
| Histone H1.4 | H14_RAT | 0 | 0 | 1 | 0 |
| Histone H4 | H4_RAT | 0 | 0 | 1 | 0 |
| Hsc70-interacting protein | F10A1_RAT | 1 | 0 | 0 | 0 |
| Hsp90 co-chaperone Cdc37 | CDC37_RAT | 0 | 0 | 0 | 1 |
| Hydroxymethylglutaryl-CoA synthase, cytoplasmic | HMCS1_RAT | 2 | 3 | 3 | 3 |
| Hypoxanthine-guanine phosphoribosyltransferase | HPRT_RAT | 0 | 1 | 0 | 0 |
| Ig kappa chain C region, A allele | KACA_RAT | 0 | 1 | 0 | 0 |
| Importin subunit beta-1 | IMB1_RAT | 0 | 0 | 2 | 3 |
| Inosine triphosphate pyrophosphatase | ITPA_RAT | 0 | 0 | 0 | 1 |
| Insulin receptor-related protein | INSRR_RAT | 0 | 1 | 0 | 0 |
| Isocitrate dehydrogenase [NAD] subunit alpha, mitochondrial | IDH3A_RAT | 0 | 1 | 0 | 0 |
| Isocitrate dehydrogenase [NAD] subunit gamma 1, mitochondrial | IDHG1_RAT | 0 | 0 | 1 | 0 |
| Isocitrate dehydrogenase [NADP] cytoplasmic | IDHC_RAT | 3 | 3 | 0 | 2 |

| | | | | | |
|---|---|---|---|---|---|
| Isocitrate dehydrogenase [NADP], mitochondrial | IDHP_RAT | 2 | 4 | 0 | 0 |
| Isoform 1 of Cytosolic acyl coenzyme A thioester hydrolase | BACH_RAT | 0 | 0 | 2 | 3 |
| Isoform 11a of Neurexin-1 | NRX1A_RAT | 0 | 0 | 2 | 2 |
| Isoform 13 of Dynamin-3 | DYN3_RAT | 0 | 0 | 0 | 2 |
| Isoform 2 of Bifunctional protein NCOAT | NCOAT_RAT | 0 | 0 | 1 | 1 |
| Isoform 2 of Clathrin coat assembly protein AP180 | AP180_RAT | 0 | 0 | 2 | 2 |
| Isoform 2 of Dynamin-1-like protein | DNM1L_RAT | 0 | 0 | 2 | 0 |
| Isoform 2 of Endoplasmin | ENPL_RAT | 0 | 0 | 1 | 1 |
| Isoform 2 of Glial fibrillary acidic protein | GFAP_RAT | 0 | 0 | 2 | 5 |
| Isoform 2 of Glutaredoxin-3 | GLRX3_RAT | 0 | 0 | 0 | 1 |
| Isoform 2 of Heterogeneous nuclear ribonucleoprotein A3 | ROA3_RAT | 0 | 0 | 0 | 1 |
| Isoform 2 of Microtubule-associated protein 6 | MAP6_RAT | 0 | 0 | 3 | 3 |
| Isoform 2 of Neuronal membrane glycoprotein M6-a | GPM6A_RAT | 1 | 2 | 1 | 0 |
| Isoform 2 of Phytanoyl-CoA hydroxylase-interacting protein-like | PHIPL_RAT | 0 | 0 | 1 | 0 |
| Isoform 2 of Protein transport protein Sec31A | SC31A_RAT | 0 | 0 | 0 | 1 |
| Isoform 2 of Reticulon-3 | RTN3_RAT | 0 | 0 | 1 | 1 |
| Isoform 2 of Sarcoplasmic/endoplasmic reticulum calcium ATPase 2 | AT2A2_RAT | 0 | 0 | 1 | 3 |
| Isoform 2 of Serine/threonine-protein phosphatase 2B catalytic subunit alpha isoform | PP2BA_RAT | 0 | 0 | 1 | 0 |
| Isoform 2 of SRC kinase signaling inhibitor 1 | SRCN1_RAT | 0 | 0 | 1 | 0 |

| | | | | | |
|---|---|---|---|---|---|
| Isoform 2 of Voltage-dependent anion-selective channel protein 3 | VDAC3_RAT | 0 | 1 | 0 | 0 |
| Isoform 2 of WD repeat-containing protein 7 | WDR7_RAT | 0 | 0 | 1 | 1 |
| Isoform 3 of Dynamin-1 | DYN1_RAT | 0 | 0 | 2 | 1 |
| Isoform 3 of NADH-cytochrome b5 reductase 3 | NB5R3_RAT | 0 | 0 | 1 | 3 |
| Isoform 3 of Neuronal-specific septin-3 | SEPT3_RAT | 0 | 0 | 2 | 1 |
| Isoform 3 of Septin-11 | SEP11_RAT | 0 | 0 | 3 | 3 |
| Isoform 4 of Dynamin-1-like protein | DNM1L_RAT | 0 | 0 | 0 | 2 |
| Isoform 4 of Heterogeneous nuclear ribonucleoprotein D0 | HNRPD_RAT | 0 | 0 | 1 | 0 |
| Isoform 4 of Protein NDRG4 | NDRG4_RAT | 0 | 0 | 0 | 1 |
| Isoform 5 of Disks large homolog 2 | DLG2_RAT | 0 | 0 | 0 | 1 |
| Isoform 5 of Dynamin-1-like protein | DNM1L_RAT | 0 | 0 | 1 | 1 |
| Isoform 5 of Synaptojanin-1 | SYNJ1_RAT | 0 | 0 | 1 | 2 |
| Isoform Cytoplasmic of Fumarate hydratase, mitochondrial | FUMH_RAT | 0 | 0 | 2 | 2 |
| Isoform Delta 5 of Calcium/calmodulin-dependent protein kinase type II subunit delta | KCC2D_RAT | 0 | 0 | 1 | 3 |
| Isoform E1 of Drebrin | DREB_RAT | 0 | 0 | 1 | 0 |
| Isoform GLAST-1A of Excitatory amino acid transporter 1 | EAA1_RAT | 0 | 0 | 2 | 3 |
| Isoform IB of Synapsin-1 | SYN1_RAT | 0 | 0 | 3 | 3 |
| Isoform II of V-type proton ATPase 116 kDa subunit a isoform 1 | VPP1_RAT | 0 | 0 | 3 | 3 |
| Isoform IIb of Synapsin-2 | SYN2_RAT | 0 | 0 | 2 | 3 |

| | | | | | |
|---|---|---|---|---|---|
| Isoform M2 of Pyruvate kinase isozymes M1/M2 | KPYM_RAT | 3 | 3 | 0 | 0 |
| Isoform Short of 14-3-3 protein beta/alpha | 1433B_RAT | 0 | 0 | 2 | 2 |
| Isoform ZA of Plasma membrane calcium-transporting ATPase 2 | AT2B2_RAT | 0 | 0 | 2 | 0 |
| Isoform ZA of Plasma membrane calcium-transporting ATPase 4 | AT2B4_RAT | 0 | 0 | 3 | 3 |
| Kinesin-1 heavy chain | KINH_RAT | 0 | 0 | 0 | 1 |
| Lactoylglutathione lyase | LGUL_RAT | 0 | 0 | 1 | 0 |
| Lamin-B1 | LMNB1_RAT | 0 | 1 | 0 | 0 |
| Large neutral amino acids transporter small subunit 1 | LAT1_RAT | 1 | 0 | 2 | 3 |
| Latexin | LXN_RAT | 0 | 0 | 0 | 1 |
| LETM1 and EF-hand domain-containing protein 1, mitochondrial | LETM1_RAT | 0 | 0 | 1 | 2 |
| Leucine-rich PPR motif-containing protein, mitochondrial | LPPRC_RAT | 1 | 3 | 0 | 0 |
| Leucine-rich repeat-containing protein 14 [LRC14_RAT] | LRC14_RAT | 0 | 0 | 0 | 2 |
| L-lactate dehydrogenase A chain | LDHA_RAT | 0 | 0 | 1 | 3 |
| L-lactate dehydrogenase B chain | LDHB_RAT | 5 | 4 | 3 | 3 |
| Lon protease homolog, mitochondrial | LONM_RAT | 0 | 0 | 1 | 1 |
| Long-chain specific acyl-CoA dehydrogenase, mitochondrial | ACADL_RAT | 3 | 1 | 0 | 0 |
| Long-chain-fatty-acid--CoA ligase 4 | ACSL4_RAT | 0 | 0 | 1 | 1 |
| Lysosome-associated membrane glycoprotein 1 | LAMP1_RAT | 2 | 3 | 1 | 1 |
| Malate dehydrogenase, cytoplasmic | MDHC_RAT | 5 | 4 | 3 | 3 |

| | | | | | |
|---|---|---|---|---|---|
| Malate dehydrogenase, mitochondrial | MDHM_RAT | 5 | 4 | 3 | 3 |
| Matrin-3 | MATR3_RAT | 0 | 0 | 1 | 2 |
| Microtubule-actin cross-linking factor 1 | MACF1_RAT | 0 | 0 | 1 | 1 |
| Microtubule-associated protein 1A | MAP1A_RAT | 2 | 2 | 1 | 2 |
| Microtubule-associated protein 1S | MAP1S_RAT | 0 | 0 | 1 | 0 |
| Microtubule-associated protein 4 | MAP4_RAT | 0 | 0 | 3 | 3 |
| Microtubule-associated protein RP/EB family member 1 | MARE1_RAT | 0 | 0 | 3 | 2 |
| Microtubule-associated protein tau | TAU_RAT | 3 | 2 | 0 | 0 |
| Mitochondrial import receptor subunit TOM70 | TOM70_RAT | 2 | 2 | 0 | 0 |
| Mitochondrial inner membrane protein (Fragment) | IMMT_RAT | 0 | 1 | 2 | 2 |
| Mitogen-activated protein kinase 1 | MK01_RAT | 0 | 0 | 2 | 3 |
| Mitogen-activated protein kinase 3 | MK03_RAT | 0 | 0 | 1 | 3 |
| Monocarboxylate transporter 1 | MOT1_RAT | 0 | 0 | 1 | 1 |
| Monocarboxylate transporter 2 | MOT2_RAT | 0 | 0 | 0 | 1 |
| Mu-crystallin homolog | CRYM_RAT | 3 | 4 | 0 | 0 |
| Multifunctional protein ADE2 | PUR6_RAT | 0 | 0 | 0 | 2 |
| Myelin transcription factor 1-like protein | MYT1L_RAT | 0 | 0 | 3 | 1 |
| Myosin light chain 1/3, skeletal muscle isoform | MYL1_RAT | 0 | 1 | 0 | 0 |
| Myosin regulatory light chain 2, skeletal muscle isoform | MLRS_RAT | 0 | 1 | 0 | 0 |
| Myosin-3 | MYH3_RAT | 0 | 1 | 0 | 0 |
| Myosin-4 | MYH4_RAT | 0 | 1 | 0 | 0 |

| | | | | | |
|---|---|---|---|---|---|
| Myosin-9 | MYH9_RAT | 1 | 0 | 0 | 0 |
| Myristoylated alanine-rich C-kinase substrate | MARCS_RAT | 0 | 2 | 0 | 0 |
| NADH dehydrogenase [ubiquinone] 1 alpha subcomplex subunit 9, mitochondrial | NDUA9_RAT | 0 | 0 | 1 | 3 |
| NADH dehydrogenase [ubiquinone] iron-sulfur protein 2, mitochondrial | NDUS2_RAT | 0 | 0 | 0 | 1 |
| NADH-ubiquinone oxidoreductase 75 kDa subunit, mitochondrial | NDUS1_RAT | 0 | 0 | 3 | 3 |
| Neogenin (Fragment) | NEO1_RAT | 0 | 0 | 0 | 1 |
| Neural cell adhesion molecule 1 | NCAM1_RAT | 3 | 4 | 0 | 0 |
| Neurochondrin | NCDN_RAT | 3 | 2 | 2 | 3 |
| Neurofilament heavy polypeptide | NFH_RAT | 0 | 0 | 0 | 2 |
| Neuromodulin | NEUM_RAT | 0 | 0 | 3 | 3 |
| Neuronal membrane glycoprotein M6-a | GPM6A_RAT | 0 | 0 | 1 | 3 |
| Non-POU domain-containing octamer-binding protein | NONO_RAT | 3 | 3 | 0 | 0 |
| Nucleolin | NUCL_RAT | 0 | 0 | 2 | 2 |
| Nucleoside diphosphate kinase B | NDKB_RAT | 0 | 1 | 0 | 0 |
| Nucleosome assembly protein 1-like 4 | NP1L4_RAT | 0 | 0 | 0 | 1 |
| Obg-like ATPase 1 | OLA1_RAT | 0 | 0 | 0 | 3 |
| Ornithine aminotransferase, mitochondrial | OAT_RAT | 0 | 0 | 1 | 2 |
| Peroxiredoxin-1 | PRDX1_RAT | 0 | 0 | 0 | 1 |
| Peroxiredoxin-2 | PRDX2_RAT | 0 | 0 | 1 | 0 |

| | | | | | |
|---|---|---|---|---|---|
| Peroxiredoxin-6 | PRDX6_RAT | 0 | 2 | 0 | 0 |
| Phosphate carrier protein, mitochondrial | MPCP_RAT | 6 | 2 | 0 | 0 |
| Phosphatidate cytidylyltransferase 2 | CDS2_RAT | 0 | 0 | 2 | 2 |
| Phosphatidylinositide phosphatase SAC1 | SAC1_RAT | 1 | 0 | 0 | 0 |
| Phosphatidylinositol transfer protein alpha isoform | PIPNA_RAT | 0 | 0 | 1 | 1 |
| Phosphoglucomutase-1 | PGM1_RAT | 2 | 3 | 0 | 0 |
| Phosphoglycerate kinase 1 | PGK1_RAT | 0 | 0 | 3 | 3 |
| Phosphoglycerate mutase 1 | PGAM1_RAT | 0 | 1 | 1 | 2 |
| Phospholipase A-2-activating protein | PLAP_RAT | 0 | 0 | 1 | 2 |
| Plasma membrane calcium-transporting ATPase 1 | AT2B1_RAT | 1 | 3 | 0 | 0 |
| Plasma membrane calcium-transporting ATPase 2 | AT2B2_RAT | 0 | 1 | 0 | 0 |
| Platelet-activating factor acetylhydrolase IB subunit alpha | LIS1_RAT | 0 | 0 | 2 | 3 |
| Polyadenylate-binding protein | PABP1_RAT | 0 | 0 | 0 | 3 |
| Polyubiquitin-B | UBB_RAT | 0 | 0 | 3 | 3 |
| Prenylcysteine oxidase | PCYOX_RAT | 0 | 0 | 2 | 2 |
| Prohibitin | PHB_RAT | 0 | 0 | 1 | 1 |
| Prohibitin-2 | PHB2_RAT | 4 | 3 | 2 | 3 |
| Prolyl endopeptidase | PPCE_RAT | 0 | 0 | 1 | 2 |
| Proteasome subunit alpha type-1 | PSA1_RAT | 0 | 0 | 0 | 1 |
| Proteasome subunit alpha type-2 | PSA2_RAT | 0 | 0 | 0 | 1 |
| Proteasome subunit alpha type-3 | PSA3_RAT | 0 | 0 | 1 | 1 |

| | | | | | |
|---|---|---|---|---|---|
| Proteasome subunit alpha type-4 | PSA4_RAT | 0 | 0 | 0 | 1 |
| Proteasome subunit alpha type-6 | PSA6_RAT | 0 | 2 | 1 | 0 |
| Proteasome subunit beta type-1 | PSB1_RAT | 0 | 1 | 0 | 1 |
| Proteasome subunit beta type-4 | PSB4_RAT | 0 | 0 | 0 | 1 |
| Proteasome subunit beta type-6 | PSB6_RAT | 0 | 0 | 0 | 1 |
| Protein disulfide-isomerase A3 | PDIA3_RAT | 4 | 3 | 3 | 3 |
| Protein disulfide-isomerase A6 | PDIA6_RAT | 0 | 0 | 3 | 3 |
| Protein disulfide-isomerase | PDIA1_RAT | 2 | 2 | 0 | 0 |
| Protein DJ-1 | PARK7_RAT | 0 | 0 | 1 | 1 |
| Protein IMPACT | IMPCT_RAT | 0 | 0 | 1 | 2 |
| Protein kinase C gamma type | KPCG_RAT | 0 | 0 | 2 | 3 |
| Protein phosphatase 1 regulatory subunit 7 | PP1R7_RAT | 0 | 0 | 0 | 2 |
| Protein phosphatase 1E | PPM1E_RAT | 0 | 0 | 1 | 0 |
| Protein phosphatase 1H | PPM1H_RAT | 0 | 0 | 1 | 0 |
| Pyruvate carboxylase, mitochondrial | PYC_RAT | 0 | 0 | 2 | 2 |
| Pyruvate dehydrogenase E1 component subunit alpha, somatic form, mitochondrial | ODPA_RAT | 2 | 3 | 2 | 2 |
| Pyruvate dehydrogenase E1 component subunit beta, mitochondrial | ODPB_RAT | 3 | 2 | 2 | 1 |
| Pyruvate kinase PKM | KPYM_RAT | 0 | 0 | 2 | 3 |
| Rab GDP dissociation inhibitor alpha | GDIA_RAT | 6 | 4 | 3 | 3 |
| Rab GDP dissociation inhibitor beta | GDIB_RAT | 4 | 4 | 2 | 3 |

| | | | | | |
|---|---|---|---|---|---|
| Ras-related C3 botulinum toxin substrate 1 | RAC1_RAT | 0 | 0 | 1 | 0 |
| Ras-related protein Rab-11A | RB11A_RAT | 0 | 0 | 1 | 0 |
| Ras-related protein Rab-11B | RB11B_RAT | 0 | 0 | 0 | 1 |
| Ras-related protein Rab-1A | RAB1A_RAT | 0 | 0 | 1 | 1 |
| Ras-related protein Rab-3A | RAB3A_RAT | 0 | 0 | 2 | 1 |
| Ras-related protein Rab-3B | RAB3B_RAT | 0 | 0 | 0 | 1 |
| Ras-related protein Rab-3C | RAB3C_RAT | 0 | 1 | 0 | 0 |
| Ras-related protein Rab-6A | RAB6A_RAT | 0 | 0 | 0 | 1 |
| Ras-related protein Rab-7a | RAB7A_RAT | 0 | 0 | 1 | 1 |
| Ras-related protein Ral-A | RALA_RAT | 0 | 1 | 1 | 1 |
| Receptor-type tyrosine-protein phosphatase alpha | PTPRA_RAT | 0 | 0 | 0 | 1 |
| Reticulon-4 | RTN4_RAT | 1 | 0 | 0 | 0 |
| Rho GDP-dissociation inhibitor 1 | GDIR1_RAT | 0 | 1 | 1 | 2 |
| Rho-related GTP-binding protein RhoB | RHOB_RAT | 0 | 1 | 0 | 0 |
| Ribose-phosphate pyrophosphokinase 1 | PRPS1_RAT | 0 | 0 | 0 | 1 |
| Sarcoplasmic/endoplasmic reticulum calcium ATPase 1 | AT2A1_RAT | 0 | 1 | 0 | 0 |
| Secernin-1 | SCRN1_RAT | 0 | 0 | 1 | 2 |
| Septin-11 | SEP11_RAT | 3 | 2 | 0 | 0 |
| Septin-2 | SEPT2_RAT | 0 | 0 | 0 | 2 |
| Septin-7 | SEPT7_RAT | 0 | 1 | 0 | 0 |
| Serine/threonine-protein kinase BRSK1 | BRSK1_RAT | 0 | 0 | 1 | 0 |

| | | | | | |
|---|---|---|---|---|---|
| Serine/threonine-protein kinase DCLK1 | DCLK1_RAT | 0 | 0 | 2 | 3 |
| Serine/threonine-protein phosphatase 2A 55 kDa regulatory subunit B alpha isoform | 2ABA_RAT | 0 | 0 | 1 | 2 |
| Serine/threonine-protein phosphatase 2A catalytic subunit alpha isoform | PP2AA_RAT | 0 | 0 | 1 | 3 |
| Serine/threonine-protein phosphatase 2A catalytic subunit beta isoform | PP2AB_RAT | 0 | 0 | 1 | 0 |
| Serine/threonine-protein phosphatase 2B catalytic subunit alpha isoform | PP2BA_RAT | 4 | 3 | 1 | 3 |
| Serine/threonine-protein phosphatase 2B catalytic subunit beta isoform | PP2BB_RAT | 0 | 0 | 2 | 2 |
| Serine/threonine-protein phosphatase PP1-alpha catalytic subunit | PP1A_RAT | 0 | 0 | 1 | 2 |
| Serine/threonine-protein phosphatase PP1-beta catalytic subunit | PP1B_RAT | 0 | 0 | 0 | 1 |
| Serine/threonine-protein phosphatase PP1-gamma catalytic subunit | PP1G_RAT | 0 | 1 | 1 | 1 |
| Serpin H1 | SERPH_RAT | 0 | 0 | 0 | 2 |
| S-formylglutathione hydrolase | ESTD_RAT | 0 | 0 | 1 | 2 |
| Short-chain specific acyl-CoA dehydrogenase, mitochondrial | ACADS_RAT | 0 | 0 | 1 | 3 |
| Sideroflexin-1 | SFXN1_RAT | 5 | 2 | 3 | 2 |
| Sideroflexin-3 | SFXN3_RAT | 0 | 0 | 2 | 3 |
| Sodium- and chloride-dependent GABA transporter 1 | SC6A1_RAT | 0 | 0 | 0 | 2 |
| Sodium/potassium-transporting ATPase subunit alpha-1 | AT1A1_RAT | 0 | 0 | 3 | 3 |

| | | | | | |
|---|---|---|---|---|---|
| Sodium/potassium-transporting ATPase subunit alpha-2 | AT1A2_RAT | 2 | 3 | 0 | 0 |
| Sodium/potassium-transporting ATPase subunit alpha-3 | AT1A3_RAT | 6 | 4 | 3 | 3 |
| Sodium/potassium-transporting ATPase subunit beta-1 | AT1B1_RAT | 5 | 4 | 3 | 3 |
| Somatotropin | SOMA_RAT | 0 | 1 | 0 | 0 |
| Sorting nexin-1 | SNX1_RAT | 1 | 0 | 0 | 0 |
| Spectrin alpha chain, non-erythrocytic 1 | SPTN1_RAT | 0 | 0 | 1 | 1 |
| Spliceosome RNA helicase Ddx39b | DX39B_RAT | 3 | 3 | 0 | 1 |
| Staphylococcal nuclease domain-containing protein 1 | SND1_RAT | 0 | 0 | 1 | 1 |
| STE20-like serine/threonine-protein kinase | SLK_RAT | 0 | 1 | 0 | 0 |
| Stress-70 protein, mitochondrial | GRP75_RAT | 3 | 3 | 0 | 0 |
| Stress-induced-phosphoprotein 1 | STIP1_RAT | 1 | 3 | 0 | 0 |
| Succinate dehydrogenase [ubiquinone] flavoprotein subunit, mitochondrial | DHSA_RAT | 2 | 2 | 0 | 0 |
| Succinate dehydrogenase [ubiquinone] iron-sulfur subunit, mitochondrial | DHSB_RAT | 0 | 1 | 0 | 0 |
| Succinyl-CoA ligase [ADP/GDP-forming] subunit alpha, mitochondrial | SUCA_RAT | 0 | 0 | 2 | 1 |
| Succinyl-CoA:3-ketoacid coenzyme A transferase 1, mitochondrial | SCOT1_RAT | 0 | 0 | 3 | 3 |
| Synapsin-1 | SYN1_RAT | 4 | 4 | 0 | 0 |
| Synapsin-2 | SYN2_RAT | 3 | 4 | 0 | 0 |
| Synaptic vesicle glycoprotein 2A | SV2A_RAT | 2 | 3 | 3 | 3 |
| Synaptic vesicle glycoprotein 2B | SV2B_RAT | 0 | 0 | 0 | 1 |

| | | | | | |
|---|---|---|---|---|---|
| Synaptic vesicle membrane protein VAT-1 homolog | VAT1_RAT | 2 | 1 | 1 | 0 |
| Synaptojanin-1 | SYNJ1_RAT | 0 | 1 | 0 | 0 |
| Synaptophysin | SYPH_RAT | 3 | 4 | 1 | 2 |
| Synaptotagmin-1 | SYT1_RAT | 5 | 4 | 3 | 3 |
| Syntaxin-1A | STX1A_RAT | 0 | 0 | 0 | 1 |
| Syntaxin-1B | STX1B_RAT | 0 | 0 | 3 | 2 |
| T-complex protein 1 subunit alpha | TCPA_RAT | 2 | 4 | 2 | 3 |
| T-complex protein 1 subunit beta | TCPB_RAT | 0 | 0 | 3 | 3 |
| T-complex protein 1 subunit delta | TCPD_RAT | 1 | 3 | 3 | 3 |
| T-complex protein 1 subunit epsilon | TCPE_RAT | 2 | 3 | 2 | 2 |
| T-complex protein 1 subunit gamma | TCPG_RAT | 5 | 3 | 2 | 3 |
| Thimet oligopeptidase | THOP1_RAT | 0 | 0 | 1 | 0 |
| Threonine--tRNA ligase, cytoplasmic | SYTC_RAT | 0 | 0 | 1 | 0 |
| Transaldolase | TALDO_RAT | 4 | 2 | 1 | 3 |
| Transcriptional activator protein Pur-alpha (Fragments) | PURA_RAT | 0 | 0 | 1 | 2 |
| Transcriptional activator protein Pur-beta | PURB_RAT | 0 | 0 | 3 | 2 |
| Transitional endoplasmic reticulum ATPase | TERA_RAT | 4 | 4 | 3 | 3 |
| Transketolase | TKT_RAT | 3 | 4 | 2 | 3 |
| Transmembrane emp24 domain-containing protein 9 | TMED9_RAT | 0 | 0 | 0 | 1 |
| Trifunctional enzyme subunit alpha, mitochondrial | ECHA_RAT | 0 | 0 | 0 | 2 |
| Trifunctional enzyme subunit beta, mitochondrial | ECHB_RAT | 1 | 3 | 0 | 1 |

| | | | | | |
|---|---|---|---|---|---|
| Triosephosphate isomerase | TPIS_RAT | 1 | 2 | 1 | 1 |
| Tripartite motif-containing protein 2 | TRIM2_RAT | 0 | 0 | 0 | 1 |
| Tripeptidyl-peptidase 2 | TPP2_RAT | 0 | 1 | 0 | 2 |
| Tropomyosin alpha-1 chain | TPM1_RAT | 0 | 1 | 0 | 0 |
| Tropomyosin alpha-3 chain | TPM3_RAT | 0 | 1 | 0 | 0 |
| Tropomyosin beta chain | TPM2_RAT | 0 | 1 | 0 | 0 |
| Troponin T, fast skeletal muscle | TNNT3_RAT | 0 | 1 | 0 | 0 |
| Tubulin alpha-1A chain | TBA1A_RAT | 0 | 0 | 5 | 3 |
| Tubulin alpha-1B chain | TBA1B_RAT | 0 | 0 | 1 | 0 |
| Tubulin alpha-4A chain | TBA4A_RAT | 0 | 0 | 1 | 3 |
| Tubulin beta-2A chain | TBB2A_RAT | 0 | 0 | 4 | 5 |
| Tubulin beta-2B chain | TBB2B_RAT | 0 | 1 | 3 | 3 |
| Tubulin beta-3 chain | TBB3_RAT | 0 | 0 | 3 | 3 |
| Tyrosine-protein kinase Fyn | FYN_RAT | 0 | 0 | 2 | 2 |
| Tyrosine--tRNA ligase, cytoplasmic | SYYC_RAT | 0 | 0 | 1 | 0 |
| Ubiquitin carboxyl-terminal hydrolase isozyme L1 | UCHL1_RAT | 1 | 2 | 1 | 1 |
| Ubiquitin thioesterase OTUB1 | OTUB1_RAT | 2 | 2 | 1 | 2 |
| Ubiquitin-like modifier-activating enzyme 1 | UBA1_RAT | 5 | 4 | 3 | 3 |
| UDP-glucose:glycoprotein glucosyltransferase 1 | UGGG1_RAT | 0 | 0 | 0 | 1 |
| Unconventional myosin-Va | MYO5A_RAT | 0 | 1 | 3 | 3 |
| Valyl-tRNA synthetase | SYVC_RAT | 3 | 3 | 0 | 0 |

| | | | | | |
|---|---|---|---|---|---|
| Vesicular glutamate transporter 1 | VGLU1_RAT | 1 | 0 | 0 | 0 |
| Vigilin | VIGLN_RAT | 0 | 1 | 0 | 0 |
| Voltage-dependent anion-selective channel protein 1 | VDAC1_RAT | 5 | 4 | 3 | 3 |
| Voltage-dependent anion-selective channel protein 2 | VDAC2_RAT | 4 | 3 | 3 | 3 |
| Voltage-dependent anion-selective channel protein 3 | VDAC3_RAT | 1 | 3 | 1 | 2 |
| Voltage-dependent calcium channel subunit alpha-2/delta-1 | CA2D1_RAT | 0 | 0 | 0 | 1 |
| V-type proton ATPase subunit B, brain isoform | VATB2_RAT | 3 | 3 | 3 | 3 |
| V-type proton ATPase subunit C 1 | VATC1_RAT | 0 | 0 | 0 | 2 |
| V-type proton ATPase subunit E 1 | VATE1_RAT | 4 | 3 | 0 | 0 |
| WD repeat-containing protein 1 | WDR1_RAT | 3 | 3 | 2 | 3 |
| Wiskott-Aldrich syndrome protein family member 1 | WASF1_RAT | 0 | 0 | 1 | 2 |
| Total: 447 | | | | | |

**Supplemental Table 2: Protein list of all identified proteins.** The columns indicate the protein name, the UniProt accession number and the number of detections of the protein in hippocampal and cortical neurons in control or NMDA-stimulated cultures.

#### SUPPLEMENTAL TABLE 3

| | **corr. p-Value** | |
|---|---|---|
| **Biological Process** | **no stimulus** | **NMDA** |
| glycolysis | 4.99E-14 | 1.62E-06 |
| tricarboxylic acid cycle | 1.29E-13 | 2.37E-06 |
| protein folding | 3.87E-12 | 6.18E-08 |

| | | |
|---|---|---|
| GTP catabolic process | 4.12E-11 | 2.91E-07 |
| ATP catabolic process | 2.50E-10 | 7.44E-08 |
| 2-oxoglutarate metabolic process | 1.03E-09 | 1.37E-05 |
| ATP biosynthetic process | 4.92E-09 | 5.80E-07 |
| carbohydrate metabolic process | 4.16E-08 | 1.60E-04 |
| synaptic transmission | 8.32E-08 | 3.72E-05 |
| oxaloacetate metabolic process | 1.69E-07 | 1.62E-03 |
| isocitrate metabolic process | 1.77E-07 | 8.57E-03 |
| NADH metabolic process | 3.31E-06 | 1.59E-02 |
| protein phosphorylation | 1.23E-04 | 3.39E-06 |
| cell cycle | 7.13E-06 | 4.49E-06 |
| cell division | 6.79E-05 | 4.90E-06 |
| pentose-phosphate shunt | 6.04E-06 | 1.87E-02 |
| response to drug | 7.16E-06 | 2.12E-04 |
| gluconeogenesis | 8.08E-06 | 5.64E-03 |
| protein export from nucleus | 4.94E-05 | 1.37E-05 |
| protein autophosphorylation | 1.94E-04 | 1.53E-05 |
| intracellular protein transport | 1.66E-05 | 1.71E-04 |
| response to organic cyclic compound | 1.69E-05 | 1.19E-03 |
| neurotransmitter transport | 1.70E-05 | 4.52E-04 |
| transport | 1.74E-05 | 8.50E-04 |
| small GTPase mediated signal transduction | 2.05E-05 | 3.30E-03 |
| response to stress | 2.23E-05 | 5.83E-05 |
| brain development | 2.47E-05 | 2.86E-05 |
| neuromuscular process controlling balance | 1.23E-04 | 2.98E-05 |
| response to toxin | 2.82E-04 | 3.63E-05 |
| cellular carbohydrate metabolic process | 3.71E-05 | 3.55E-02 |
| response to hypoxia | 4.13E-05 | 2.17E-04 |

| | | |
|---|---|---|
| protein transport | 4.94E-05 | 1.30E-04 |
| glutathione metabolic process | 5.17E-05 | 1.62E-03 |
| endocytosis | 6.39E-05 | 6.01E-04 |
| translational elongation | 7.97E-05 | 4.74E-03 |
| ionotropic glutamate receptor signaling pathway | 1.05E-04 | 1.18E-03 |
| metabolic process | 1.12E-04 | 9.81E-03 |
| neuron projection development | 1.14E-04 | 4.18E-03 |
| regulation of protein localization | 1.15E-04 | 3.04E-04 |
| axonogenesis | 1.23E-04 | 1.49E-04 |
| nervous system development | 1.31E-04 | 2.73E-03 |
| protein dephosphorylation | 1.35E-04 | 4.40E-03 |
| aging | 4.36E-04 | 2.03E-04 |
| vesicle-mediated transport | 2.16E-04 | 2.15E-03 |
| NAD metabolic process | 2.18E-04 | |
| regulation of the force of heart contraction | 6.85E-04 | 2.22E-04 |
| response to reactive oxygen species | 6.85E-04 | 2.22E-04 |
| muscle filament sliding | 4.94E-04 | 2.36E-04 |
| regulation of long-term neuronal synaptic plasticity | 3.18E-04 | 9.27E-03 |
| microtubule-based process | 1.36E-03 | 5.54E-04 |
| cellular response to growth factor stimulus | 2.22E-03 | 5.69E-04 |
| neural retina development | 1.21E-03 | 6.00E-04 |
| regulation of stress-activated MAPK cascade | 1.21E-03 | 6.00E-04 |
| microtubule-based movement | 6.44E-04 | 6.21E-03 |
| proteolysis | 6.89E-04 | 2.92E-02 |
| response to cocaine | 7.02E-04 | 1.74E-03 |
| regulation of heart contraction | 1.82E-03 | 7.24E-04 |
| response to nutrient | 1.04E-03 | 8.30E-04 |
| acetyl-CoA biosynthetic process from pyruvate | 8.36E-04 | |

| | | |
|---|---|---|
| pyrimidine base catabolic process | 8.36E-04 | |
| negative regulation of microtubule depolymerization | 8.54E-04 | 3.68E-03 |
| calcium ion transport | 8.66E-04 | 4.63E-03 |
| protein polymerization | 8.88E-04 | 2.06E-03 |
| regulation of exocytosis | 1.01E-03 | 4.24E-03 |
| organ regeneration | 4.14E-03 | 1.13E-03 |
| response to cytokine stimulus | 1.13E-03 | 1.21E-03 |
| actin filament capping | 2.21E-03 | 1.18E-03 |
| citrate metabolic process | 1.21E-03 | 1.08E-02 |
| malate metabolic process | 1.21E-03 | |
| regulation of protein transport | 1.21E-03 | |
| response to electrical stimulus | 2.86E-03 | 1.21E-03 |
| response to morphine | 2.86E-03 | 1.21E-03 |
| protein homotetramerization | 3.44E-03 | 1.23E-03 |
| response to heat | 1.24E-03 | 1.69E-03 |
| cellular calcium ion homeostasis | 1.25E-03 | 1.70E-03 |
| response to ethanol | 1.84E-03 | 1.33E-03 |
| calcium ion transmembrane transport | 3.32E-03 | 1.33E-03 |
| positive regulation of anti-apoptosis | 1.34E-03 | 2.84E-03 |
| mitosis | 5.86E-03 | 1.46E-03 |
| caveolin-mediated endocytosis | 2.10E-03 | 1.48E-03 |
| regulation of interferon-gamma-mediated signaling pathway | 2.10E-03 | 1.48E-03 |
| regulation of type I interferon-mediated signaling pathway | 2.10E-03 | 1.48E-03 |
| muscle contraction | 4.10E-03 | 1.54E-03 |
| phosphorylation | 4.10E-03 | 1.54E-03 |
| regulation of multicellular organism growth | 4.10E-03 | 1.54E-03 |
| response to estrogen stimulus | 1.66E-03 | 1.55E-03 |
| myoblast fusion | 3.42E-03 | 1.62E-03 |

| | | |
|---|---|---|
| purine nucleotide biosynthetic process | 3.42E-03 | 1.62E-03 |
| liver development | 1.82E-03 | 1.69E-03 |
| cation transport | 6.40E-03 | 1.70E-03 |
| anti-apoptosis | 1.73E-03 | 1.85E-02 |
| multicellular organismal development | 1.80E-03 | 1.74E-03 |
| bone resorption | 1.82E-03 | 4.54E-02 |
| cholesterol biosynthetic process | 1.82E-03 | 4.54E-02 |
| glucose metabolic process | 1.83E-03 | 3.62E-03 |
| learning or memory | 1.83E-03 | 3.62E-03 |
| regulation of cell shape | 1.84E-03 | |
| insulin-like growth factor receptor signaling pathway | 4.24E-03 | 1.86E-03 |
| response to growth hormone stimulus | 4.24E-03 | 1.86E-03 |
| response to calcium ion | 1.99E-03 | 2.71E-03 |
| learning | 6.21E-03 | 2.06E-03 |
| microtubule cytoskeleton organization | 2.08E-03 | 4.25E-03 |
| fatty acid metabolic process | 2.09E-03 | 2.80E-03 |
| cellular response to oxidative stress | 2.10E-03 | 6.79E-03 |
| aldehyde catabolic process | 2.10E-03 | 4.56E-02 |
| glyceraldehyde-3-phosphate metabolic process | 2.10E-03 | |
| glyoxylate cycle | 2.10E-03 | 4.56E-02 |
| ribose phosphate biosynthetic process | 2.10E-03 | 4.56E-02 |
| uropod organization | 2.10E-03 | 4.56E-02 |
| actin cytoskeleton organization | 2.11E-03 | 3.93E-02 |
| ATP metabolic process | 2.13E-03 | |
| electron transport chain | 2.23E-03 | 4.51E-03 |
| glycogen metabolic process | 2.50E-03 | 8.53E-03 |
| intracellular protein kinase cascade | 2.68E-03 | 1.73E-02 |
| labyrinthine layer blood vessel development | 6.19E-03 | 2.79E-03 |

| | | |
|---|---|---|
| MAPK import into nucleus | 4.91E-03 | 2.84E-03 |
| chaperone-mediated protein complex assembly | 4.91E-03 | 2.84E-03 |
| ketone body catabolic process | 4.91E-03 | 2.84E-03 |
| pentose-phosphate shunt, oxidative branch | 4.91E-03 | 2.84E-03 |
| positive regulation of dopamine metabolic process | 4.91E-03 | 2.84E-03 |
| regulation of muscle contraction | 7.37E-03 | 3.12E-03 |
| forebrain development | 1.15E-02 | 3.12E-03 |
| oxidation-reduction process | 3.31E-03 | 5.54E-03 |
| neuron migration | 3.46E-03 | 4.18E-03 |
| sarcomere organization | 8.98E-03 | 4.24E-03 |
| microtubule organizing center organization | 7.91E-03 | 4.38E-03 |
| protein import into mitochondrial outer membrane | 7.91E-03 | 4.38E-03 |
| protein localization to microtubule | 7.91E-03 | 4.38E-03 |
| regulation of ATPase activity | 7.91E-03 | 4.38E-03 |
| regulation of Golgi inheritance | 7.91E-03 | 4.38E-03 |
| regulation of early endosome to late endosome transport | 7.91E-03 | 4.38E-03 |
| regulation of protein export from nucleus | 7.91E-03 | 4.38E-03 |
| regulation of striated muscle contraction | 7.91E-03 | 4.38E-03 |
| cell adhesion | 1.41E-02 | 4.43E-03 |
| response to organic substance | 6.34E-03 | 4.51E-03 |
| endoplasmic reticulum tubular network organization | 4.91E-03 | |
| gamma-aminobutyric acid metabolic process | 4.91E-03 | |
| response to food | 1.15E-02 | 4.96E-03 |
| response to hormone stimulus | 5.16E-03 | 5.41E-03 |
| protein heterooligomerization | 2.14E-02 | 5.83E-03 |
| response to estradiol stimulus | 8.37E-03 | 5.84E-03 |
| chaperone mediated protein folding requiring cofactor | 6.19E-03 | |
| response to amphetamine | 6.21E-03 | 1.60E-02 |

| | | |
|---|---|---|
| Rab protein signal transduction | 1.14E-02 | 6.22E-03 |
| cell morphogenesis involved in differentiation | 1.14E-02 | 6.22E-03 |
| fructose 1,6-bisphosphate metabolic process | 1.14E-02 | 6.22E-03 |
| negative regulation of necrotic cell death | 1.14E-02 | 6.22E-03 |
| succinyl-CoA metabolic process | 1.14E-02 | 6.22E-03 |
| negative regulation of apoptotic process | 8.34E-03 | 6.23E-03 |
| epithelial to mesenchymal transition | 1.50E-02 | 6.26E-03 |
| response to organic nitrogen | 6.40E-03 | 1.01E-02 |
| ventricular cardiac muscle tissue morphogenesis | 1.62E-02 | 6.79E-03 |
| peptidyl-serine phosphorylation | 2.13E-02 | 7.18E-03 |
| regulation of protein catabolic process | 7.37E-03 | 3.55E-02 |
| lipid metabolic process | 1.10E-02 | 7.80E-03 |
| cellular response to inorganic substance | 7.91E-03 | |
| glutamate biosynthetic process | 7.91E-03 | |
| neurotransmitter biosynthetic process | 7.91E-03 | |
| pentose-phosphate shunt. non-oxidative branch | 7.91E-03 | |
| adult locomotory behavior | 8.09E-03 | 2.00E-02 |
| visual learning | 8.25E-03 | 1.91E-02 |
| cell differentiation | 8.34E-03 | 3.66E-02 |
| lipopolysaccharide-mediated signaling pathway | 2.03E-02 | 8.53E-03 |
| apical protein localization | 1.54E-02 | 8.57E-03 |
| synaptic vesicle maturation | 1.54E-02 | 8.57E-03 |
| synaptic vesicle transport | 1.54E-02 | 8.57E-03 |
| calcium ion-dependent exocytosis | 8.60E-03 | 3.90E-02 |
| establishment of cell polarity | 8.60E-03 | 3.90E-02 |
| ATP hydrolysis coupled proton transport | 8.82E-03 | |
| cytokinesis | 8.82E-03 | |
| regulation of cell migration | 8.82E-03 | |

| | | |
|---|---|---|
| cell migration | 8.85E-03 | |
| protein homooligomerization | 8.85E-03 | |
| hydrogen peroxide catabolic process | 8.98E-03 | 3.22E-02 |
| response to vitamin E | 8.98E-03 | 3.22E-02 |

**Supplemental Table 3.** Filtered Genecodis results: Single enrichment analyses were performed with Genecodis on the SNO-proteome data of the NMDA stimulated and unstimulated rats. The resulting lists were merged via their Gene Ontology Terms (biological processes). A cut-off for the adjusted p-values was set at 0.01 and p-values cells were filled grey, if and only if the p-values were at least $1x10^{-1}$ apart, marking the group with potentially more proteins dedicated to the biological process.
